# Neurodynamics and connectivity during facial fear perception: The role of threat exposure and signal congruity

**DOI:** 10.1101/149112

**Authors:** Cody Cushing, Hee Yeon Im, Reginald B. Adams, Noreen Ward, Daniel N. Albohn, Troy G. Steiner, Kestutis Kveraga

## Abstract

Fearful faces convey threat cues whose meaning is contextualized by eye gaze: While averted gaze is congruent with facial fear (both signal avoidance), direct gaze is incongruent with it, as direct gaze signals approach. We have previously shown using fMRI that the amygdala is engaged more strongly by fear with averted gaze, which has been found to be processed more efficiently, during brief exposures. However, the amygdala also responds more to fear with direct gaze during longer exposures. Here we examined previously unexplored brain oscillatory responses to characterize the neurodynamics and connectivity during brief (∼250 ms) and longer (∼883 ms) exposures of fearful faces with direct or averted eye gaze. We replicated the exposure time by gaze direction interaction in fMRI (N=23), and observed greater early phase locking to averted-gaze fear (congruent threat signal) with MEG (N=60) in a network of face processing regions, with both brief and longer exposures. Phase locking to direct-gaze fear (incongruent threat signal) then increased significantly for brief exposures at ∼350 ms, and at ∼700 ms for longer exposures. Our results characterize the stages of congruent and incongruent facial threat signal processing and show that stimulus exposure strongly affects the onset and duration of these stages.

## Introduction

When we look at a face, we can glean a wealth of information, such as age, sex, health, affective state, and attentional focus. The latter two signals are typically, but not exclusively, carried by emotional expression and eye gaze direction, respectively. Depending on the emotional expression and gaze, we can recognize how happy, angry, or fearful a person is, and infer the source or target of that emotion [1]. In an initial examination of the interaction between eye gaze and facial emotion, Adams and colleagues introduced the “shared signal hypothesis” [2–4]. This hypothesis predicts that when paired, cues relevant to threat expression, and sharing a congruent underlying signal value, should facilitate the processing efficiency of that expression. In support of this hypothesis, using speeded reaction time tasks and self-reported intensity of emotion perceived, Adams and Kleck [3, 4] found that direct gaze facilitated processing efficiency and accuracy, and increased the perceived emotional intensity of approach-oriented emotions (e.g., anger and joy). Conversely, averted gaze facilitated perception of avoidance-oriented emotions (e.g., fear and sadness). Several other groups have now also found similar results, including a replication by Sander et al. [5] using dynamic threat displays, another using a diffusion model of decision making and reaction time [6], and another examining effects on reflexive orienting to threat [7]. Perhaps the most compelling behavioral replication of this effect was a study by Milders and colleagues [8], who found that direct-gaze anger and averted-gaze fear were detected more readily in an attentional blink paradigm compared to averted-gaze anger and direct-gaze fear, suggesting that congruent pairings (i.e., shared signals) of gaze and emotion attract more preconscious attentional awareness. Similar interaction effects have been found at the neural level as well, including in several fMRI studies looking at amygdala responses to different gaze directions by threat displays, including our own (see too [9–11]).

Some have also suggested that gaze influences the ambiguity surrounding the source of threat. Whalen and colleagues, for instance, hypothesized that amygdala activation is directly proportional to the amount of ambiguity that surrounds the source of a perceived threat [12], which suggests that direct-gaze fear is a more ambiguous signal than direct-gaze anger. Anger signals both the source of the threat and where it is directed. In the case of fear, the observer knows there is a threat, but direct eye gaze does not indicate the source of the threat - unless it is the observer. Thus, averted eye gaze is more informative in resolving the source of threat for fear, and direct gaze is more informative for anger. In both of these accounts of threat-related ambiguity (shared signals and source of threat detection), direct-gaze fear is considered a more ambiguous combination of cues than averted-gaze fear.

Initial efforts to study the neural underpinnings of the perception of these compound threat cues revealed greater amygdala activation in response to incongruent compound threat cues, specifically fearful faces with a direct gaze and anger faces with averted gaze [13, 14]. Some follow-up studies, however, have found the opposite interaction: fear with averted eye gaze evoked higher amygdala activation [9, 10]. To address this, Adams et al. [15] proposed that presentation duration might help explain these differences. We hypothesized that brief presentations trigger more *reflexive* processing, which is thought to be preferentially tuned to congruent threat cues (averted-gaze fear), and longer presentations engage more r*eflective* processing of the less salient, incongruent threat cues (direct-gaze fear, see e.g. [16–18] for discussions of *reflexive* vs. *reflective* processing). Indeed, previous studies finding stronger amygdala activation in response to averted-gaze fear used relatively brief stimulus exposure times (e.g. [9] used 300 ms stimulus durations), whereas those reporting higher amygdala activation in response to direct-gaze fear had longer stimulus exposure times (e.g, [14, 19] used 2s and 1s stimulus durations, respectively). Adams et al. [15] put this hypothesis to the test in the context of three studies using fMRI, with varying presentation parameters during a constant 1.5s trial. In a direct comparison, this work revealed that amygdala responses were enhanced for fearful faces coupled with averted gaze, when rapidly presented (300ms), and to fearful faces coupled with direct eye gaze, when presented for a sustained duration (1s).

## The Current Work

The primary goal of the present study was to elucidate the fine-grained neural dynamics of this previously observed response by replicating and extending this work using magnetoencephalography (MEG) to record neural activity in response to brief (250 ms) and longer (883 ms) presentations of fearful faces with direct and averted gaze. MEG allows us to elucidate not only the temporal evolution of neural activity, but also frequency-specific oscillatory activity in response to the stimulus, including highly temporally resolved interregional connectivity patterns during perception of these compound threat cues from the face. We utilized source localization to obtain good spatial resolution to identify the temporally sensitive contributions of key brain regions in the extended face processing network: the Fusiform Face Area (FFA), Periamygdaloid Cortex (PAC), posterior superior temporal sulcus (pSTS), and orbitofrontal cortex (OFC), as well as the earliest cortical visual region V1. These regions are well known to be involved in either face perception and social communication or gaze perception, if not both (e.g., [20–24]. STS in general has been shown to be sensitive to gaze [25], as has the fusiform gyrus [26]. In addition, both have been shown to be sensitive to facial expression [27]. The STS and OFC are also implicated as nodes in the proposed “social brain” consisting of amygdala, OFC, and STS [28]. The posterior portion of STS also has been implicated as specializing in inferring intentionality from social cues [29, 30]. Whether or not amygdala activity can be source localized from MEG data is actively debated in the MEG literature. However, there is accumulating evidence now that MEG activity can indeed be localized to the subcortical nuclei of the amygdala [31–35], but supporting or advancing this claim is not the goal of our manuscript. Periamygdaloid cortex (PAC) is heavily involved in conveying inputs and outputs of the deeper amygdala nuclei, the contralateral PAC, as well many other cortical regions [36, 37]. While we cannot be certain which of the amygdala nuclei the activity is coming from (a situation similar to all but the highest resolution fMRI studies), given the reliable activation of the amygdala in all of the previous studies using this paradigm it is probable that at least some of the activity may arise in the subcortical nuclei of the amygdala.

To examine the frequency-specific responses in our ROIs and assess functional connectivity between them, we computed phase-locking estimates in and between the evoked responses of these regions. Phase-locking (in which the magnitude of oscillatory activity is normalized) was chosen over power analyses (a magnitude-dependent measure of synchronized neuronal firing) due to the fact that phase locking is by nature more informative of the timing of activity within a region as well as the timing of functional connectivity between regions [38]. Phase locking is also thought to be a more trustworthy measure of higher-level functions [39]. Oscillations in α-band (8-13 Hz) have been implicated as being involved in task selection or disengagement from the task [40–43] while β-band (13-30 Hz) activity is indicative of active cognitive processing [44–47]. By examining activity in these bands both locally and between regions, we sought to elucidate the neurodynamics underlying the differential processing of threat cues portrayed by direct and averted gaze on a fearful face observed by Adams et al. [15] when exposure duration is manipulated.

We were primarily interested in characterizing how exposure duration within a constant time frame influences threat processing as a function of threat cue congruity associated with direct or averted gaze, previously found to evoke differential activation using fMRI [15]. An adaptive response to threat cues would be a combination of reflexive and reflective processes that enables appropriate and timely responses to clear threat signals (e.g., fleeing an attacker) while also inhibiting context-inappropriate or maladaptive responses to more ambiguous threat cues (e.g., fleeing from someone who is seeking help). Temporally, initial reflexive processing has been found to begin as early as 50-100 ms, becoming fully engaged by 300 ms, while intentional responses have been found as early as 500-700 ms [48–51]. Thus, we predicted that the initial response would be stronger to averted-gaze fear within this timeframe, in agreement with the findings of Milders et al. [8], which found that the congruent averted-gaze fear signal attracted more preconscious attention than direct-gaze fear. On the other hand, we predicted that direct-gaze fear would evoke a stronger secondary response indicative of reflective analysis to resolve the conflicting signals in the incongruent cue. Moreover, we hypothesized that this late-arising reflective response would be greater during longer stimulus exposures, as suggested by the fMRI results [14, 15, 19, 52]

Finally, we also wanted to test whether the effects reported in van der Zwaag et al. [52] and Adams et al. [15] can still be evoked with rapid switching of conditions in an event-related design (both direct and averted-gaze fear faces, and brief/longer exposure durations), rather than state-dependent, which could be the case for previous work employing block designs. Therefore, we employed a rapid event-related design in MEG and also scanned a separate cohort in fMRI using an identical paradigm. An additional motivation was to compare the periamygdaloid complex activation in MEG to the fMRI results to test whether the PAC activity we found in MEG is at least broadly comparable to the BOLD activity in fMRI. While BOLD activity is unable to capture the fine-grained temporal dynamics present in the MEG signal, the overall activation differences might be comparable [53].

## Methods & Materials

### Experiment 1

#### Participants

Total 28 undergraduate students (22 females, mean age (s.d.)=18.75 (1.73)) participated in Experiment 1. All the participants had normal color vision and normal or corrected-to-normal visual acuity. Their informed written consent was obtained according to the procedures of the Institutional Review Board at the Pennsylvania State University. The participants received a course credit for their participation. This research was performed in accordance with the guidelines and regulations set forth in the Declaration of Helsinki, and was approved by the Institutional Review Board of the Pennsylvania State University.

#### Stimuli

The face stimuli in this experiment were identical to those used in the three studies reported in Adams et al. [15], with eight models (four female) from the Pictures of Facial Affect [54] and eight models (four female) from the NimStim Emotional Face Stimuli database [55], all displaying a fearful expression with either a direct gaze or averted gaze (Figure 1A). Each model had 6 stimuli unique to their identity: one with direct gaze, one with leftward averted gaze, one with rightward averted gaze, and then the mirror image of each of these iterations. This resulted in 96 unique face stimuli used in the experiment. Images of ordinary household objects were also used to ensure observer attention and required a response. Stimuli were presented with Psychtoolbox [56, 57] in Matlab (Mathworks Inc., Natick, MA). Stimuli were projected onto a screen at the head of the bore and viewed via an angled mirror attached to the head coil subtending approximately 5.72° horizontally and 7.77° vertically of visual angle.

**Figure 1.**
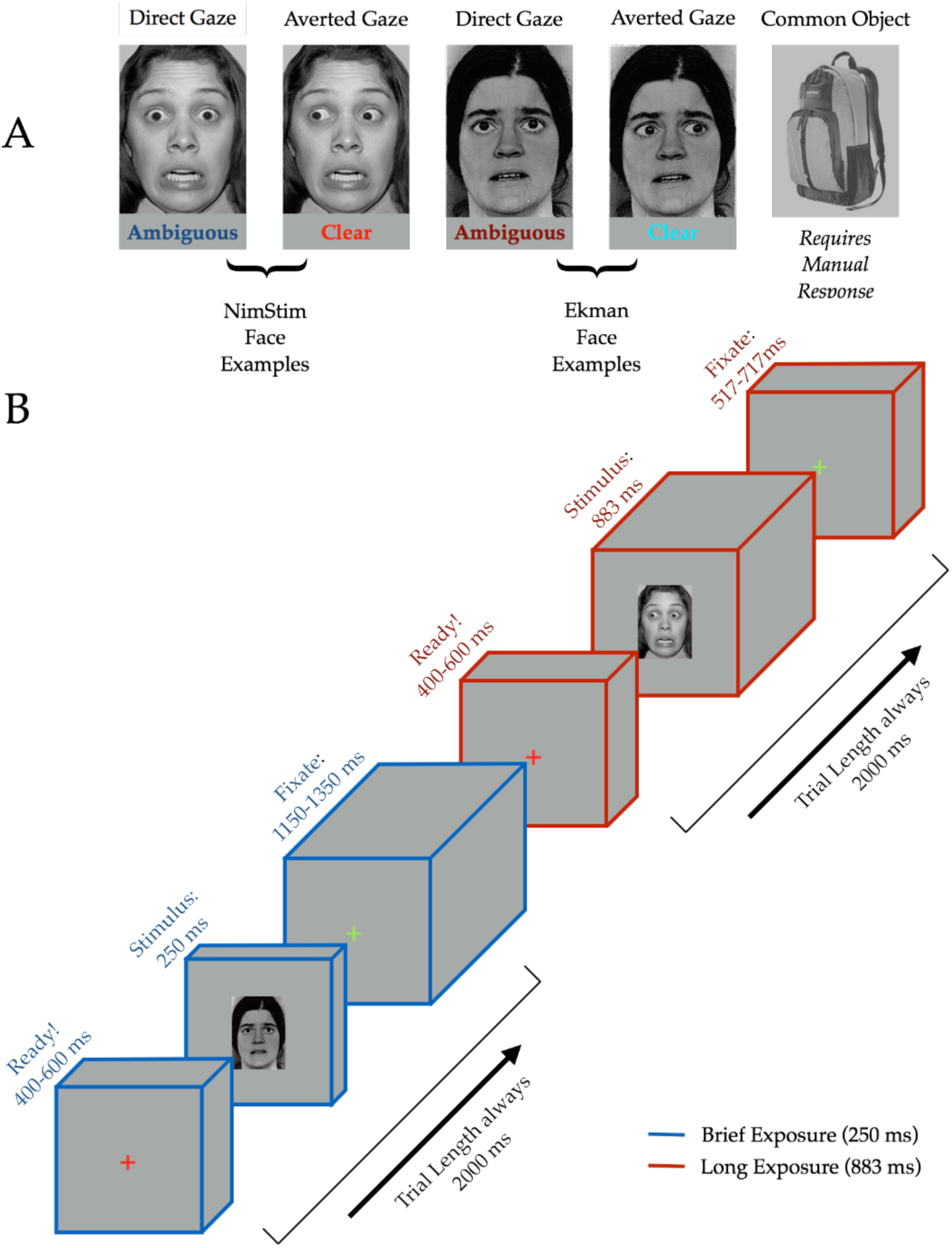
Stimuli and Task Design. **(A)** Examples of threat cue and object stimuli. All faces were taken from the NimStim or Ekman databases (examples of both shown) and displayed a fearful expression with either direct or averted gaze. Participants were instructed to make a button response if the stimulus was a non-face object, ensuring attentive viewing of all faces. **(B)** Sequence depicting one each of both brief (blue cuboids) and long (red cuboids) exposure trial types. Trials are always 2000 ms: 400-600ms of a red fixation cross signifying trial start, followed by 250 ms or 883 ms of stimulus (corresponding to trial type), concluding with either 1150-1350ms or 517-717 ms of a green fixation cross, dependent upon trial type. End-trial jitter is inversely timed with pre-trial jitter such that trial lengths are kept a constant 2000 ms.

#### Task Design

We employed an event-related design while keeping presentation parameters as similar to Adams et al. [15] as possible within such a design. Each participant viewed 320 trials over the four runs that were randomly, evenly split into brief (15 frames at a refresh rate of 16.67 ms totaling 250 ms) and longer stimulus (53 frames at a refresh rate of 16.67 ms totaling 883 ms) durations to get 160 trials for each presentation duration (128 of which were faces). Each trial lasted 2 seconds, beginning with a randomized 400-600 ms of attending to a central red fixation cross. The stimulus (either a face or an object) was then presented for either 250 ms or 883 ms, dependent upon trial type, followed by a green fixation cross for the remainder of the trial (ranging from 517-1350 ms) before switching back to red to signify the start of a new trial (Figure 1B). Participants were instructed only to make a manual response via a button box if the stimulus was a non-face object. This task would allow us to ensure that participants pay attention to faces without explicitly labeling the emotion displayed, as previous studies have shown that the act of emotion labeling changes neural responsivity to the emotional expression [58].

#### fMRI Data Acquisition and Analysis

fMRI images of brain activity were acquired using a 3T scanner (Siemens Magnetom Prisma) located at The Pennsylvania State University Social, Life, and Engineering Sciences Imaging Center. High-resolution anatomical MRI data were acquired using T1-weighted images for the reconstruction of each subject’s cortical surface (TR = 2300 ms, TE = 2.28 ms, flip angle = 8°, FoV = 256 x 256 mm^2^, slice thickness = 1 mm, sagittal orientation). The functional scans were acquired using gradient-echo EPI with a TR of 2000 ms, TE of 28ms, flip angle of 52° and 64 interleaved slices (3 x 3 x 2 mm resolution). Scanning parameters were optimized by manual shimming of the gradients to fit the brain anatomy of each subject, and tilting the slice prescription anteriorly 20-30° up from the AC-PC line as described in the previous studies [59, 60] to improve signal and minimize susceptibility artifacts in the brain regions susceptible to signal dropout, such as OFC [61]. We acquired 456 functional volumes per subject in four functional runs, and the sequence of trials was optimized for hemodynamic response estimation efficiency using *optseq2* software (https://surfer.nmr.mgh.harvard.edu/optseq/).

The acquired fMRI data were analyzed using SPM8 (Wellcome Department of Cognitive Neurology, http://www.fil.ion.ucl.ac.uk/spm/software/spm8/). The functional images were corrected for differences in slice timing, realigned, corrected for movement-related artifacts, coregistered with each participant’s anatomical data, normalized to the Montreal Neurological Institute template, and spatially smoothed using an isotropic 8-mm full width half-maximum (FWHM) Gaussian kernel. ArtRepair software was used to correct for excessive movement (http://spnl.stanford.edu/tools/ArtRepair/ArtRepair.htm), and outliers due to movement or signal from preprocessed files, using thresholds of 3 s.d. from the mean, 0.75 mm for translation and 0.02 radians rotation, were removed from the data sets [62]. Data of five participants among the 28 participants were unusable with ArtRepair identifying > 75% of scans as outliers. Therefore, only the remaining 23 participants were included for further fMRI analyses.

Subject-specific contrasts were estimated using a fixed-effects model. These contrast images were used to obtain subject-specific estimates for each effect. For group analysis, these estimates were then entered into a second-level analysis treating participants as a random effect, using one-sample t-tests at each voxel. We computed contrasts between brief and longer exposures within each Threat type (i.e., brief averted-gaze fear vs. longer averted-gaze fear, brief direct-gaze fear vs. longer direct-gaze fear) and between averted-gaze fear and direct-gaze fear within each exposure duration, separately. The full lists of these whole brain activations are shown in Table 1 and Table 2, thresholded at t > 3.0 (p < 0.003) and a minimal cluster size of 5 voxels.

**Table 1.** Regions showing increased activation in Experiment 1 in brief-minus longer-exposure, and longer-minus brief-exposure contrasts for averted-gaze fear faces (clear threat) and direct-gaze fear faces (ambiguous threat) (height: *t*(22)>3.0, *p*<0.0033, extent=5 voxels).

|  |  | MNI Coordinates |  |  |  |
| --- | --- | --- | --- | --- | --- |
| Region Label |  |  |  |  |  |
| *Averted gaze* |  |  |  |  |  |
| *Brief - Long* | t-value | Extent | x | y | z |
| L Brainstem | 4.797 | 160 | -3 | -10 | -26 |
| R Brainstem | 3.738 | 38 | 9 | -28 | -46 |
| L Parahippocampal cortex | 4.289 | 160 | -24 | -7 | -30 |
| R Parahippocampal cortex | 3.055 | 8 | -60 | -7 | -34 |
| R Amygdala | 4.574 | 30 | 27 | 2 | -26 |
| L Frontal cortex (BA8, Frontal eye fields) | 4.498 | 31 | -24 | 17 | 42 |
|  | 3.443 | 20 | -30 | 23 | 54 |
| L Posterior superior temporal sulcus | 4.397 | 61 | -42 | -49 | 14 |
| R Posterior superior temporal sulcus | 3.098 | 62 | 43 | -60 | 13 |
| L Premotor cortex (BA6) | 4.232 | 74 | -21 | -10 | 58 |
| L Premotor cortex | 3.591 | 53 | -33 | 2 | 30 |
| L Inferior parietal lobule | 4.162 | 233 | -30 | -55 | 38 |
| L Cerebellum | 4.139 | 17 | -24 | -43 | -38 |
| R Cerebellum | 3.808 | 14 | 12 | -64 | -28 |
| R Inferior frontal gyrus | 4.099 | 127 | 42 | 11 | 30 |
| L Pallidum | 3.757 | 38 | -9 | -1 | -2 |
| L Superior temporal cortex | 3.625 | 8 | -39 | -28 | -2 |
| R Middle temporal cortex | 3.433 | 62 | 69 | -28 | -20 |
| L Precentral cortex | 3.249 | 16 | -51 | 8 | 38 |
| R Supplementary motor area | 3.085 | 23 | 21 | 11 | 54 |
| R Dorsolateral prefrontal cortex | 3.073 | 7 | 45 | 35 | 38 |
| *Long - Brief* |  |  |  |  |  |
| L Occipital cortex (BA 18) | 6.314 | 1181 | -18 | -100 | 6 |
| R Occipital cortex (BA 17) | 6.259 | 1181 | 15 | -97 | 2 |
| R Occipital cortex (BA 18) | 3.871 | 1181 | 33 | -94 | 14 |
| *Direct gaze* |  |  |  |  |  |
| *Brief-Long* |  |  |  |  |  |
| None |  |  |  |  |  |
| *Long - Brief* |  |  |  |  |  |
| L Occipital cortex (BA 18) | 3.248 | 12901 | 36 | -82 | -4 |
| R Occipital cortex (BA 18) | 9.223 | 12901 | 21 | -94 | -8 |
| L Occipital cortex (BA 19) | 7.340 | 12901 | -27 | -79 | -10 |
| L Fusiform cortex | 7.340 | 6496 | -33 | -79 | -10 |
| R Fusiform cortex | 6.290 | 8734 | 36 | -52 | -12 |
| L Brainstem | 6.727 | 12901 | -6 | -34 | -24 |
| R Brainstem | 3.698 | 29 | 6 | -34 | -50 |
| L Posterior cingulate cortex | 8.005 | 120 | -12 | -43 | 26 |
| R Posterior cingulate cortex | 3.862 | 63 | 12 | -34 | 22 |
| R Parahippocampal cortex | 5.315 | 97 | 27 | 2 | -36 |
| R Middle cingulate cortex | 4.784 | 137 | 9 | -4 | 40 |
| L Temporal pole | 4.724 | 100 | -39 | 14 | -32 |
| R Temporal pole | 3.740 | 26 | 30 | 14 | -34 |
| R Lateral orbitofrontal cortex | 4.131 | 18 | 51 | 39 | -13 |
| R Middle temporal cortex | 4.072 | 133 | 57 | -7 | -20 |
| R Precuneus | 3.036 | 133 | 15 | -49 | 78 |
| L Medial orbitofrontal cortex | 3.929 | 53 | -18 | 12 | -17 |
| R Postcentral area | 3.908 | 87 | 48 | -25 | 40 |
| L Lateral orbitofrontal cortex | 3.890 | 75 | -54 | 29 | -10 |
| L Insula | 3.710 | 36 | -36 | -22 | 22 |
| L Superior frontal gyrus (BA6) | 3.631 | 28 | -45 | 5 | 32 |
| L Inferior frontal gyrus | 3.496 | 15 | -48 | 14 | 10 |
| L Inferior temporal cortex | 3.466 | 25 | -51 | -10 | -32 |
|  | 3.463 | 28 | -36 | -7 | -38 |
|  | 3.196 | 6 | -54 | -25 | -22 |
| L Anterior superior temporal cortex | 3.391 | 5 | -51 | -4 | -6 |
| L Posterior superior temporal cortex | 3.331 | 51 | -60 | -34 | 0 |
| L Cerebellum | 3.232 | 10 | -24 | -55 | -44 |
| L Anterior orbitofrontal cortex | 3.096 | 8 | -27 | 38 | -16 |
| L Amygdala | 3.090 | 13 | -15 | 2 | -22 |
| R Amygdala | 3.076 | 18 | 21 | -1 | -22 |

**Table 2.**
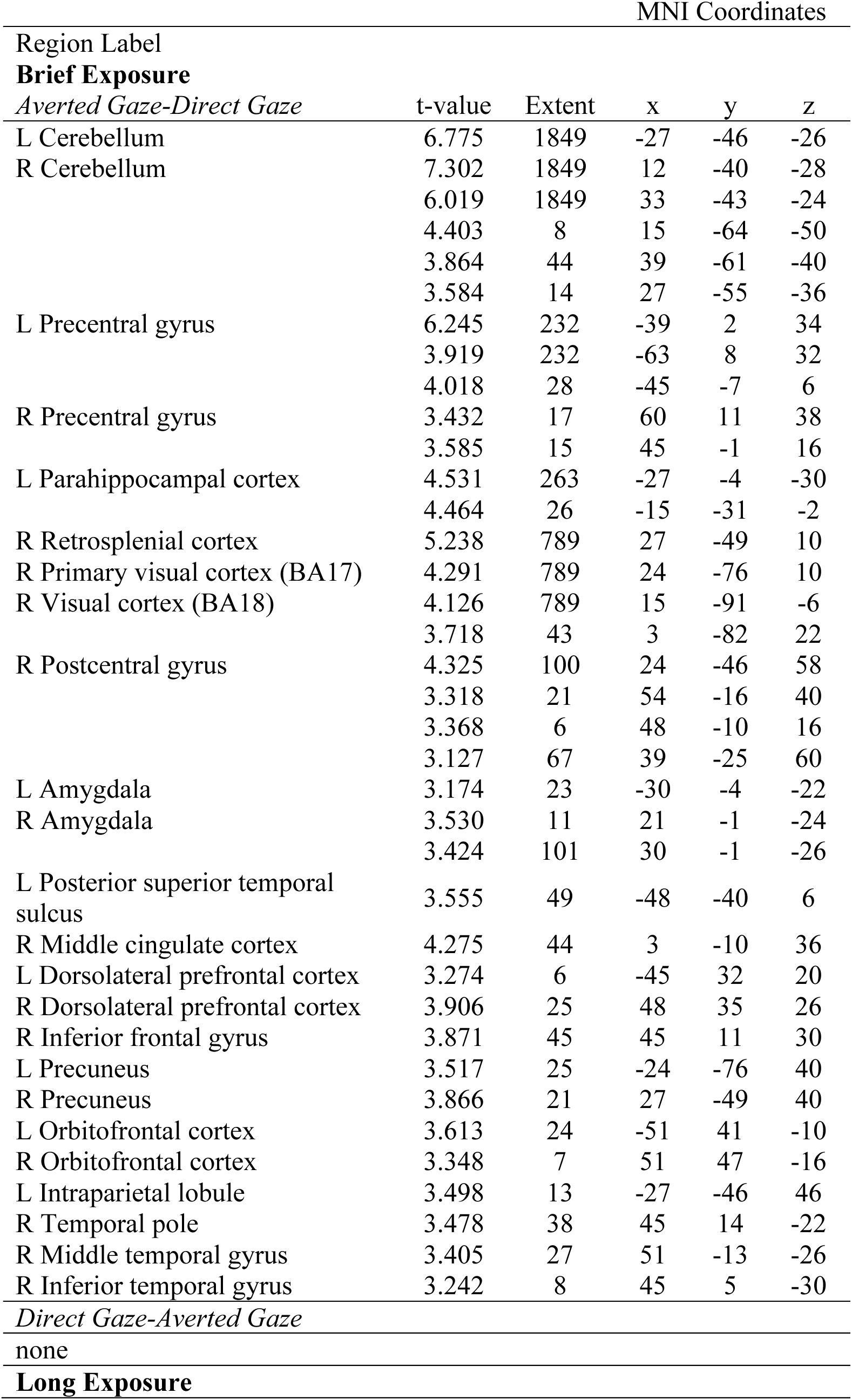
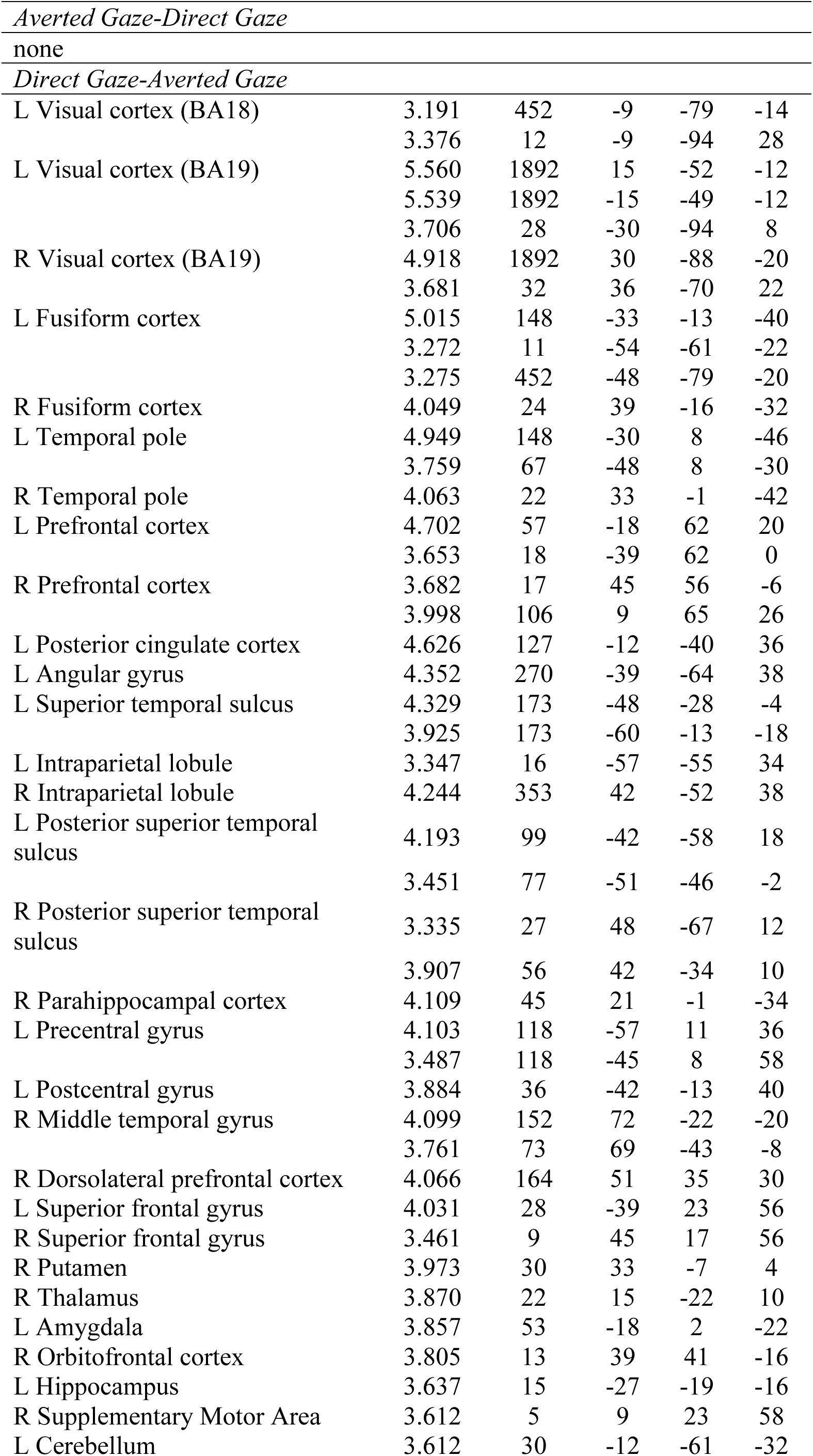
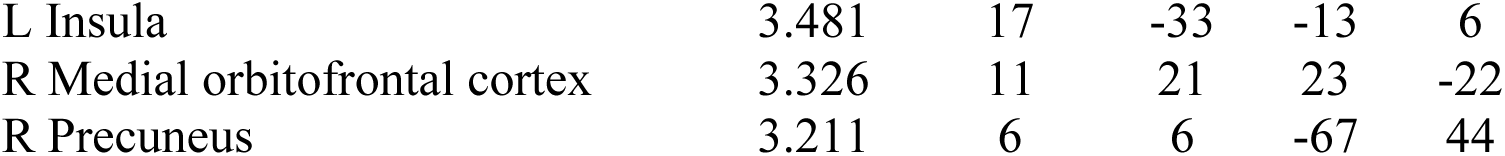
Regions showing increased activation in Experiment 1 in averted-gaze (clear threat) minus direct-gaze (ambiguous threat) and direct-gaze minus averted-gaze contrasts during brief and long exposures (height: *t*(22)>3.0, *p*<0.0033, extent=5 voxels).

For the regions of interest (ROI) analyses, we extracted the BOLD activity from our *a priori* ROIs: the amygdala, FFA, OFC, and pSTS. We defined another contrast between all the visual stimulation trials (the brief and longer exposures of averted-gaze fear and direct-gaze fear) vs. Null trials. From this contrast, we extracted the percent signal change in our ROIs for all the four conditions using the rfxplot toolbox (http://rfxplot.sourceforge.net) for SPM. We identified the [x y z] coordinate for each of our ROIs (the MNI coordinates are shown in Figure 2) then defined a 6mm sphere around it. The coordinates for Amygdala, FFA, and OFC were determined based on the previous research that reported the involvement of these regions in processing emotional facial expression (e.g., [14, 15]). Using the rfxplot toolbox in SPM8, we extracted all the voxels from each individual participant’s functional data within that sphere. The extracted percent signal change for each of the four trial conditions was subjected to a two-way repeated-measures ANOVA with the factors of Exposure duration (2 levels: brief (250 ms) vs. longer (883 ms)) and Threat type (2 levels: averted-gaze fear vs. direct-gaze fear), separately for each of the ROIs. In addition, due to our previous findings of left amygdala being specifically sensitive to direct-gaze fear (incongruent threat cue) during the longer exposures [15], we performed a planned comparison for left hemisphere ROIs of longer exposure direct-gaze fear to all other conditions.

**Figure 2.**
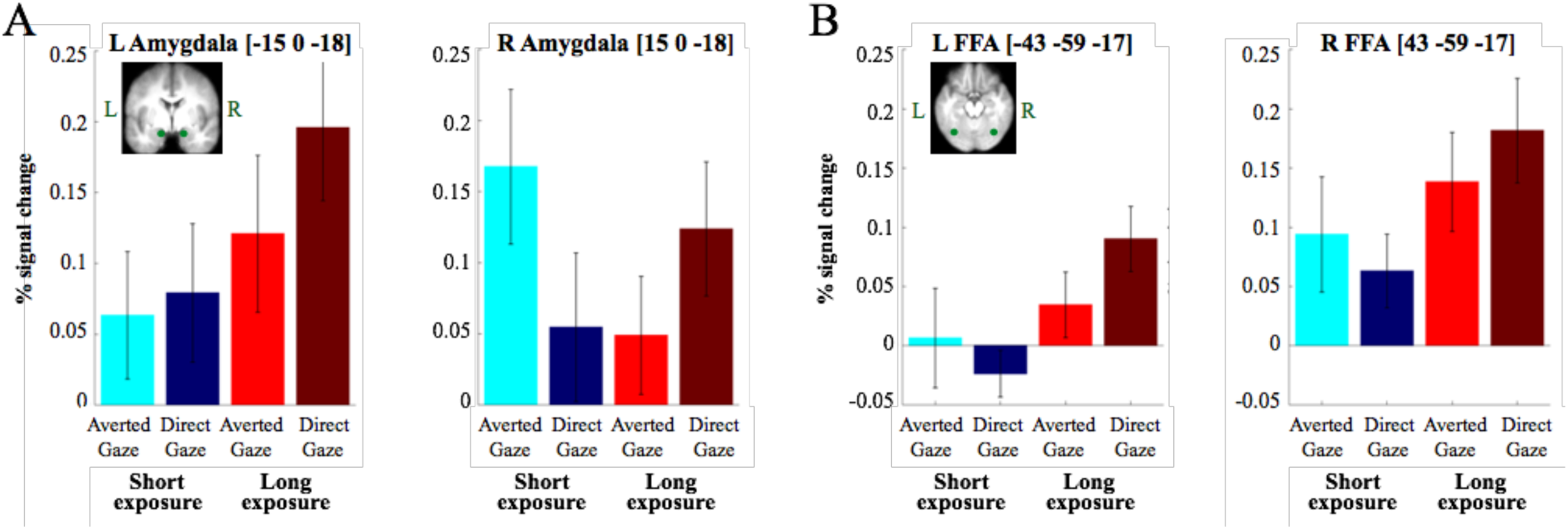
Results from Experiment 1 (fMRI study). **(A)** Shows the percent change in the BOLD signal in bilateral amygdala (left=left hemisphere, right=right hemisphere) in response to brief (blue colors) and long (red colors) to clear threat cues (brighter colors) and ambiguous threat cues (darker colors). **(B)** Shows the percent change in the BOLD signal in bilateral Fusiform Face Area (FFA) (left=left hemisphere, right=right hemisphere) in response to brief (blue colors) and long (red colors) to clear threat cues (brighter colors) and ambiguous threat cues (darker colors). Note for both (A) and (B), that a clear threat cue is represented by a fearful face with averted eye-gaze and an ambiguous threat cue is represented by a fearful face with direct eye-gaze.

### Experiment 2

#### Participants

Sixty participants (40 females, mean age (s.d.) = 26.6 (6.9)) with normal or corrected to normal vision completed the study for monetary compensation ($50). Potential subjects were screened via a questionnaire to make sure they were eligible for MEG recording and subsequent MRI structural scans, and had no history of mental illness or use of psychoactive medication. Their informed written consent was obtained according to the protocol approved by the Institutional Review Board of MGH. This research was performed in accordance with the guidelines and regulations set forth in the Declaration of Helsinki, and was approved by the Institutional Review Board of the Massachusetts General Hospital.

#### Stimuli & Task Design

Stimuli and task design were identical to Experiment 1 with the exception of breaks between runs in the MEG being self-paced until the participant was ready to continue. Stimuli were rear-projected onto a translucent screen placed 160 cm from the seated participant to create a 61.5 cm x 38.5 cm display. Stimuli measured 14.1 x 19.2 cm of subtending about 5.1° visual angle horizontally and 6.9° vertically.

#### MEG Acquisition

Magnetoencephalogram recordings were obtained with a 306-channel Neuromag Vectorview whole-head system (Elekta Neuromag) with 204 planar gradiometers and 102 magnetometers enclosed in a magnetically shielded room with a shielding factor of 250,000 at 1 Hz (ImedcoAG). Four head position indicator (HPI) electrodes were affixed asymmetrically to each participant’s forehead and the mastoid processes to monitor head position in the dewar at the beginning of the recording session. Digitizer data were collected for each participant’s head on a Polhemus FastTrack 3D system within a head-coordinate frame defined by anatomical landmarks (left preauricular area, right preauricular area, and the nasion). HPI positions were marked within this frame, and 150-200 points on the scalp and the face were entered for co-registration with structural MRIs of the subject. Eye movements and blinks were monitored via 4 EOG electrodes: 2 vertical electrodes on the left eye (one placed just above the eyebrow, the other on the upper cheekbone just below the eye), and 2 horizontal electrodes (placed on either side of the head between the eye and hairline). Cardiac activity was recorded via ECG using electrodes placed on the left and right chest (2 total). All data from MEG sensors and EOG and ECG electrodes were sampled at 600.615 Hz and were band-pass filtered at 0.1-200 Hz. Recordings were stored for offline analysis.

#### Data Pre-processing and Averaging

All recordings were pre-processed and averaged using a combination of the MNE analysis package [63] as well as MNE-Python [64] and custom scripts in Python and Matlab. Signal-space projection was applied to the recordings in order to remove noise from external sources [65, 66]. Sensors that were visibly noisy during the recording were noted by the researchers and excluded from analysis. For time course analysis, a low-pass filter of 40 Hz was applied, and recordings were epoched from 200 ms before stimulus onset to 1300 ms post-stimulus. For time-frequency analysis, no filter was applied to the data, and recordings were epoched from 500 ms before stimulus onset until 1440 ms post-stimulus onset. Rejection parameters were set at 4,000 fT/cm for gradiometers, 4000 fT for magnetometers, and 800 uV for EOG. Any epoch where any of these limits were exceeded was excluded from further analysis. A further data quality inspection was performed during preprocessing and any noisy or flat channels that were not picked up during the recording, but resulted in the rejection of 20% or more of epochs were excluded from analysis to prevent unnecessary epoch rejection. No participants were excluded due to excessive trial rejection. The lowest number of trials entered into the analyses for a condition for any subject was 51 trials. There were no significant differences between conditions for trial count (all *p*’s>0.86). We excluded trials where object images were presented from MEG analyses as they were of no experimental interest.

#### Source Localization

A structural MRI for each participant was acquired on a 1.5T Siemens Avanto 32-channel “TIM” system. A single compartment boundary-element model was fitted to the intracranial volume of the MRI data in the form of a triangular mesh isomorphic to an icosahedron recursively divided 5 times. This model was implemented in a surface-based forward solution to restrict the sources of the MEG signal to the vertices of this triangular mesh (source space) fitted to each individual’s inflated cortical surface reconstructed using the Freesurfer analysis package [67]. Sources closer than 5mm to the inner skull surface were omitted from the forward solution. The MRI-head coordinate transformation for each subject was supplied to the forward model by aligning the digitizer data obtained in the original recording session (see *MEG Acquisition*) with a high-resolution head surface tessellation constructed from the MRI data. The inverse operator was prepared with a loose orientation constraint (LOC) parameter of 0.2 in order to improve localization accuracy [68]. A depth-weighting coefficient of 0.8 was also set for the inverse operator to lessen the tendency of minimum-norm estimates to be localized to superficial currents in place of deep sources. Only gradiometers were used in the depth weighting process. Both gradiometers and magnetometers were used to source localize the data. MEG data were source localized onto the whole brain using a lambda2 regularization parameter based on Signal-to-Noise Ratio (SNR) equal to 1/(SNR^2^). Evoked cortical activation was quantified spatiotemporally by taking only the radial component from a 3-orientation source [x y z] at each vertex in the form of dynamic statistical parametric maps (dSPMs) based on an inverse solution regularized with an SNR of 3. These are a statistical representation of significant activity from each source per time point calculated by noise-normalization on the estimated current amplitude (MNE) of a given source according to noise covariance between sensors calculated during a baseline period of 200 ms pre-stimulus [69]. The noise covariance estimation model was selected automatically according to rank for each participant [70]. To analyze the spectral content of the neural response, MEG data were source localized on a trial-by-trial basis using the minimum-norm estimate (MNE) method with the same inverse operator regularized with an SNR of 1 due to the indiscernibility of signal and noise at the single trial level.

#### ROI Selection and Definition

Regions of interest (ROIs) were chosen based on their established roles in early visual processing, face perception, threat detection, and emotional processing: early visual cortex (V1), fusiform face area (FFA), posterior superior temporal sulcus (pSTS), periamygdaloid cortex (PAC), orbitofrontal cortex (OFC). For the MEG data analyses, the ROIs (‘labels’ in the terminology of the *mne_analyze* software) were functionally derived in each individual’s anatomical space within *a priori* anatomical constraints (automatic cortical parcellations) produced with the Freesurfer analysis package [67], except for the PAC and pSTS as explained below. The functional label within the anatomical parcellation was derived from averaged activity from all conditions, so that the activity was independent of trial type. This enabled us to account for intersubject variability in regions like FFA, pSTS, and OFC. Functional labels were generated within the anatomical parcellation corresponding to the ROI by isolating the source-space vertex with the highest activation within the anatomical constraints as well as neighboring vertices in the source-space (also within the anatomical constraints) that reach at least 60% of the maximum activation (in dSPM values). Since no suitable *a priori* parcellation of the periamygdaloid cortex was available, a posteriori anatomical constraints were imposed in the form of user-drawn ROIs on the *fsaverage* inflated surface corresponding to the cortex surrounding the amygdala (periamygdaloid cortex). The drawing of the PAC labels was tracked by linking the drawn points to be displayed on the *fsaverage* MRI volume in *tkmedit* to ensure that only the cortical surface around the amygdalae was included in the label. These anatomical constraints were then morphed to each individual’s inflated surface and used to generate functional PAC ROIs according to the preceding procedure. A similar method was used to obtain the posterior portion of STS as the *a priori* parcellation generated by Freesurfer extended beyond the true pSTS on many subjects’ cortical surfaces to inferior sulci. User-marked constraints on the *fsaverage* inflated surface were marked around STS and tracked in the *fsaverage* MRI volume. The same morphing procedure from above was used, and then the label was split into thirds. The most posterior third was then taken as each individual’s pSTS to be used as the anatomical constraint when generating the functional pSTS labels.

#### Time Course Analysis

Time courses were produced for each ROI by averaging the activity from source-space vertices that fell within the label marked on the individual’s inflated cortical surface to be submitted to statistical analysis. The individual average activity was then further averaged across subjects in order to visualize the grand average.

#### Phase-locking Analysis

Using modified scripts from the MNE-Python package [64], the Phase-Locking Factor (PLF) across trials was calculated for each ROI, and Phase-Locking Value (PLV) was calculated to assess functional connectivity between two ROIs. The PLF is a number between 0 and 1 (1 representing perfect synchrony) that represents a magnitude-normalized measure of the phase angle consistency across trials for a particular time-point at a particular frequency [71]. This number was obtained by source localizing each epoch into source space using the Minimum-Norm Estimate (MNE) method with the sign of the signal preserved. Source-space MNE epochs were subjected to spectral decomposition at each time point for each frequency of interest, using a continuous wavelet transformation with a family of complex morlet wavelets containing a number of cycles equal to f/7, where f denotes the frequency of interest. This keeps the time window at each frequency identical resulting in stable temporal resolution across frequency ranges. We analyzed frequencies from 8 Hz (representing the lower limit of the α-band) to 30 Hz (representing the upper limit of the β-band). To make these results easier to interpret, and in attempt to localize effects away from spectral leakage inherent in such transformations, PLFs were analyzed in separate frequency ranges: α (8-13 Hz), β (13-30 Hz). Similarly, inter-regional connectivity was assessed with PLVs, also a magnitude-normalized measure of phase-angle consistency across trials between 2 ROIs. This was calculated with the same parameters on the same frequencies as above (8-30Hz) and analyzed by frequency band.

#### Statistical Analyses

As this work is a direct extension of Adams et al. [15], we approached this work with particular effects whose temporal properties we sought to investigate. Specifically, we knew that averted-gaze fear elicits stronger BOLD activity relative to direct-gaze fear during brief presentations and the opposite is true of longer presentations. Consequently, we performed non-parametric comparisons based on *t*-tests comparing direct vs. averted-gaze fear faces within each presentation duration to describe the temporal evolution of these opposing sensitivities. Additionally, to verify our presentation duration manipulations were functioning as intended (i.e., to perform a reality check), we compared brief vs. longer presentations within each gaze direction, also using non-parametric comparisons based on *t*-tests. All statistics were computed using non-parametric cluster-level permutation tests based on 5000 permutations with a critical alpha-value of 0.05, following Maris and Oostenveld [72]. Cluster mass was determined by summing t-values within the cluster rather than counting significant pixels/time-points. Reported *p*-values are Monte Carlo *p-*values comparing the observed cluster to a null distribution comprising the largest cluster yielded by permuted data sets. That is, the reported *p-*value is the percentage of permuted data sets that yielded a larger cluster than the actual observed cluster (e.g., *p*=0.05 means 250 out 5000 permuted data sets yielded a larger cluster than the observed cluster).

##### Time Domain

Statistical analyses in the time domain were performed by subtracting each participant’s evoked response for 2 conditions of interest to create a contrast wave. Null distributions were built by means of a sign-flip permutation based on a one-sample t-test. Observed clusters of significant time-points whose masses were exceeded by 5 percent of or fewer clusters from the null distribution were considered significant.

##### Time-Frequency Domain

Phase-locking maps for each participant (2-dimensional images where the y-axis represents frequency and the x-axis represents time with pixel values corresponding to phase-locking factors or values) were smoothed via a Gaussian image filter with a kernel size of 5 and a sigma of 2 before being submitted to permutations and statistical analysis. Permutations were performed by shuffling condition labels for each participant, such that the condition label of each participant’s phase-locking map was randomized but no one subject ended up with both phase-locking maps (the unit of observation in this case) falling under the same condition. Both the observed and permuted statistical maps were thresholded at an alpha-level of 0.05 with 59 degrees of freedom in order to identify clusters. Observed clusters of contiguous supra-threshold time-frequency points whose masses were exceeded by 5 percent or less of clusters from the null distribution were considered significant.

##### Data Availability

All data and code used to perform analyses reported herein are available from the corresponding author at reasonable request.

## Results

### fMRI Results

Our main interest was in the amygdala responses to averted-gaze vs. direct-gaze fear (congruent vs. incongruent threat cues) and their interactions with brief vs. longer stimulus exposures. Figure 2A shows the percentage of BOLD signal change from the baseline for each of the four trial conditions (the brief and longer exposures of averted-gaze fear and direct-gaze fear) in the left and right amygdala. A two-way repeated measures ANOVA with the factors of the Exposure duration (2 levels: brief [250 ms] vs. longer [883 ms]) and the Threat type (2 levels: averted-gaze fear vs. direct-gaze fear) showed a significant main effect of exposure duration in the left amygdala (*F*(1,22) = 7.19, *p* < 0.015), such that the left amygdala (Figure 2A, left panel) showed greater activation for longer exposure than for brief exposure of threat cues (Figure 2A, left panel). However, neither the main effect of Threat type (*F*(1,22) = 1.564, *p* = 0.224) nor the interaction between the Exposure duration and the Threat type (*F*(1,22) = 0.496, *p* = 0.489) were statistically significant. Based on our *a priori* hypothesis that the left amygdala would be more involved in the sustained processing of the incongruent threat cue (the longer exposure of direct-gaze fear), we conducted a planned comparison to compare the longer exposure of direct-gaze fear with any other conditions and observed a marginally significant effect (*t*(22) = 1.70, *p* = 0.093).

In the right amygdala (Figure 2A, right panel), both main effects of the Exposure duration (*F*(1,22) = 0.295, *p* = 0.593) and of the Threat type (*F*(1,22) = 0.208, *p* = 0.652) were not significant. However, the predicted interaction between Exposure duration and Threat type was significant (*F*(1,22) = 7.19, *p* < 0.015). Specifically, the right amygdala responded more strongly to the averted-gaze fear when the exposure was brief and to direct-gaze fear when the exposure was longer. These results are consistent with previous findings that indicate possible amygdala lateralization in orientation and evaluation of facial threat cues [73].

In the left FFA (Figure 2B, left panel), we found significantly greater activation for longer exposure over brief exposure, confirmed by a significant main effect of the Exposure duration (*F*(1,22) = 5.491, *p* < 0.03). Although a main effect of Threat type (*F*(1,22) = 0.284, *p* = 0.600) and the interaction between Exposure duration and Threat type (*F*(1,22) = 1.766, *p* = 0.198) were not statistically significant, the same planned comparison as in the left amygdala (i.e., the longer exposure of direct-gaze fear vs. other conditions) showed a significant effect (*t*(22) = 2.076, *p* < 0.05). In the right FFA (Figure 2B, right panel), the main effect of Exposure duration was also significant (*F*(1,22) = 6.620, *p* < 0.02). Furthermore, we observed a similar activation pattern in the right FFA as in the right amygdala: The brief exposure of averted-gaze fear and the longer exposure of direct-gaze fear elicited stronger right FFA responses than their counterparts (brief exposure of direct-gaze fear and longer exposure of averted-gaze fear, respectively, see Fig. 2B, right panel). However, the interaction between Exposure duration and Threat type (*F*(1,22) = 1.221, *p* = 0.281) was not significant nor was the main effect of Threat type (*F*(1,22) = 0.029, *p* = 0.867). Thus, the left and right FFA responses somewhat resembled those in the left and right amygdala, but were weaker: Greater activation was observed for the longer exposure of direct-gaze fear in the left FFA and greater activation for the brief exposure of averted-gaze fear and the longer exposure of direct-gaze fear in the right FFA. It is also worth noting that the right FFA (Figure 2B, right panel) showed greater levels of activation than the left FFA (Figure 2B, left panel) in all the conditions, consistent with previous findings on the right hemisphere dominance in function of FFA [74–77]. To confirm this statistically, we conducted a separate three-way repeated measures ANOVA with additional factor of Hemisphere (left FFA versus right FFA) along with the Exposure duration and Threat type, and found a significant main effect of Hemisphere (*F*(1,22) = 7.465, *p* < 0.015).

Consistent with the ROI results, the statistical analyses on the whole brain using a univariate GLM approach showed the greater right amygdala activation for the brief exposure of averted-gaze fear, compared to the longer exposure, and the greater bilateral amygdala for the longer exposure of direct-gaze fear, compared to the brief exposure (Table 1). As shown in Table 1, we also observed that the other ROIs including the OFC, Fusiform area, V1, and the pSTS were differentially activated as a function of the Exposure duration by Threat type interaction, with greater responses for the brief exposure of averted-gaze fear and the longer exposure of direct-gaze fear. Supporting this finding at the whole brain level, there is unanimously stronger activity to averted-gaze fear relative to direct-gaze fear during brief exposures and to direct-gaze fear compared to averted-gaze fear during longer exposures (see Table 2).

### MEG Results

#### Effects of Exposure Duration: Time Courses

The time courses of activation within our ROIs revealed an effect of stimulus exposure duration for both averted-gaze fear and direct-gaze fear. The time courses can be seen in Figure 3, and a comprehensive list of the timing and significance of these effects can be found in Table 3. In V1 and FFA, we found significantly increased activation for brief compared to longer exposures only in the right hemisphere and only for averted-gaze fear for V1 (*p*=0.02) and direct-gaze fear for FFA (*p*=0.04). Conversely, we observed significantly increased activation from longer presentations over brief presentations bilaterally for both gaze directions in V1 and FFA. Bilateral V1 and right FFA were the only regions in which we observed significantly greater activity from the longer-exposure condition, while the stimulus was still on the screen, compared to the brief-exposure condition after stimulus offset (Fig. 3). In pSTS we observed a similar temporal progression, but found significant differences in evoked activity between exposure durations only during the viewing of direct-gaze fear faces (see Table 3 for timing and significance). PAC and OFC bilaterally demonstrated a distinctly different temporal progression of activity compared to the other regions (Figure 3). This was characterized in the brief-exposure condition by a rise in activity starting around 500 ms, hundreds of milliseconds after the stimulus offset at 250 ms. This increase superseded activity in the longer-exposure condition at around 600 ms, even though the stimulus was still being displayed (see Figure 3 and Table 3 for the timing and statistics of significant differences). This activity in response to the brief-exposure stimulus remained stronger until around 1000 ms (750 ms after stimulus offset) when a similar pattern began to emerge from the longer exposure, evoked by the removal of the threat cue. Thus, in pSTS, PAC, and OFC, the activity after the stimulus offset (post-250 ms) in the brief-exposure condition was higher, and showed a different pattern, than activity in the longer-exposure condition in which the stimulus was still present (250-883 ms).

**Figure 3.**
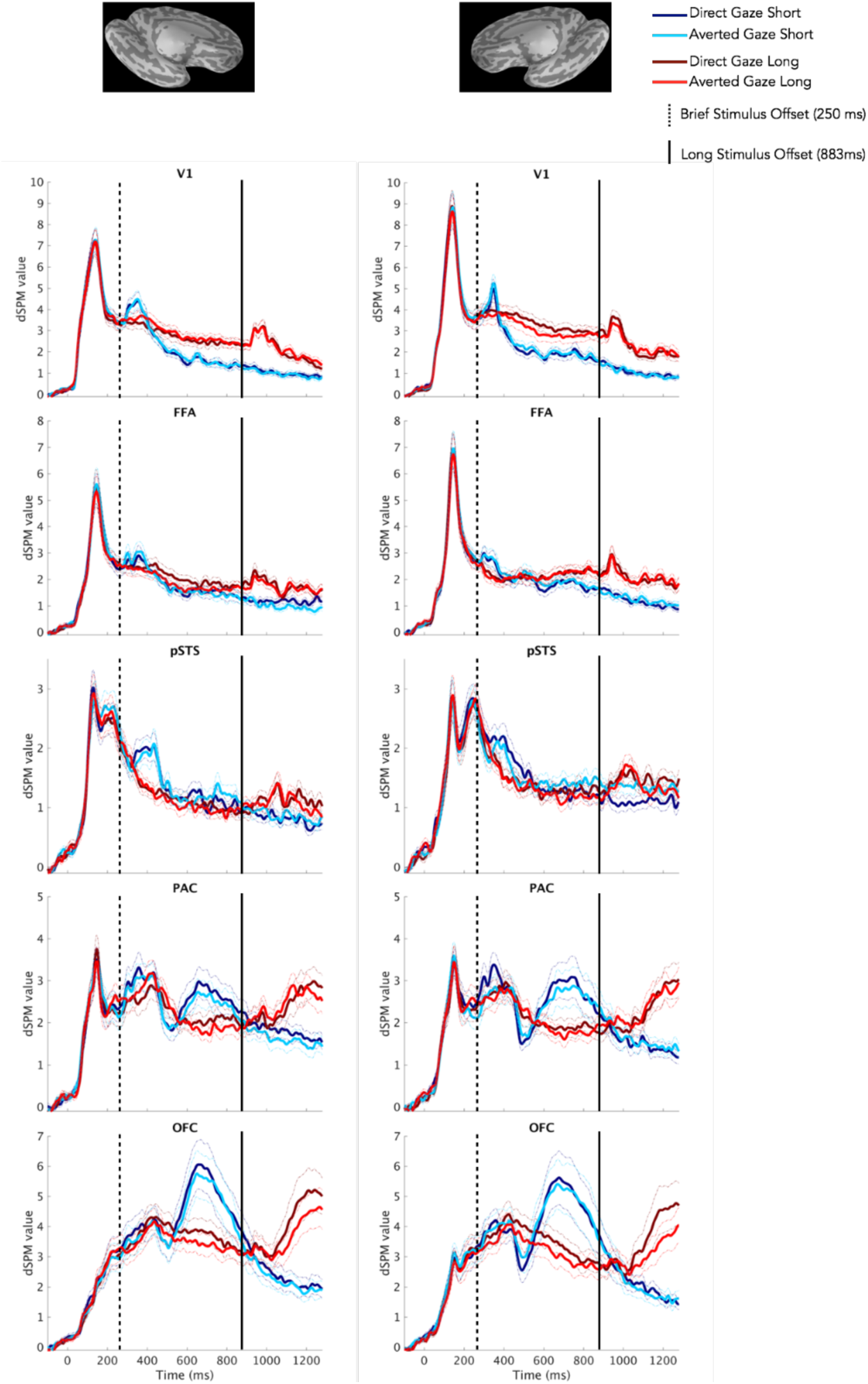
Time courses of activation within Regions of Interest (ROIs). Time courses depicting bilateral (Left=Left Hemisphere, Right=Right Hemisphere) activation (measured by dSPM values) for both exposure durations (blue=short, red=long) and both threat cue types (brighter shades=averted gaze [“clear” threat], darker shades=direct gaze [“ambiguous” threat]) in our Regions of Interest (ROIs). Standard Error from the Mean (SEM) is represented by dashed lines of the same color above and below the respective time series. The dashed black line indicates stimulus offset for brief exposures (250 ms) while the solid black line indicates stimulus offset for long exposures (883 ms).

**Table 3.** Timing of significant differences in activation within ROIs between brief (250 ms) and long (883 ms) exposures (reported *p*-values are non-parametric and corrected based on cluster permutations thresholded at *p*<0.05 uncorrected).

| ROI | LH |  | RH |  |
| --- | --- | --- | --- | --- |
| Averted Gaze |  |  |  |  |
| Brief minus Long | Time (ms) | p-value | Time (ms) | p-value |
| V1 | n.s. | n.s. | 150-220<br>320-365 | 0.01<br>0.02 |
| FFA | n.s. | n.s. | n.s | n.s |
| pSTS | n.s | n.s | n.s | n.s |
| PAC | n.s. | n.s. | 620-820 | 0.001 |
| OFC | 600-800 | 0.02 | 600-820 | 0.01 |
| Long minus Brief |  |  |  |  |
| V1 | 500-1300 | 0.0004 | 420-1300 | 0.0002 |
| FFA | 900-1300 | 0.0004 | 800-1300 | 0.0002 |
| pSTS | n.s | n.s | n.s | n.s |
| PAC | 1020-1300 | 0.002 | 1100-1300 | 0.0006 |
| OFC | 1075-1300 | 0.004 | 1100-1300 | 0.002 |
| Direct Gaze |  |  |  |  |
| Brief minus Long |  |  |  |  |
| V1 | n.s. | ns. | n.s. | n.s |
| FFA | n.s. | n.s. | 300-380 | 0.04 |
| PSTS | 380-480 | 0.05 | n.s. | n.s. |
| PAC | n.s. | n.s. | 620-780 | 0.03 |
| OFC | n.s. | n.s. | 620-800 | 0.02 |
| Long minus Brief |  |  |  |  |
| V1 | 500-1300 | 0.0002 | 400-1300 | 0.0002 |
| FFA | 900-1080 | 0.02 | 800-1300 | 0.0002 |
| pSTS | 975-1300 | 0.003 | 950-1100 | 0.04 |
| PAC | 1100-1300 | 0.03 | 1100-1300 | 0.001 |
| OFC | 1050-1300 | 0.01 | 1050-1300 | 0.0008 |

In summary, effects of exposure duration dominated the activity in all ROIs we examined. The nature of these effects seemed to depend on the role of the region in the processing hierarchy, as PAC and OFC, showed strongest responses after stimulus offset. To examine these effects in greater detail, we performed phase-locking analyses in all the ROIs for frequencies in the α and β bands.

#### Effects of Exposure Duration: Phase locking

Examination of phase synchrony across trials in our ROIs also yielded powerful effects of stimulus exposure duration for both averted-gaze and direct-gaze fear (Figure 4). See Table 4 and Table 5 for the exact timing and significance of these effects in the α and β frequency ranges, respectively. Similar to the activity we observed in the time courses, there were effects of exposure duration in every ROI examined. Phase-locking exposure effects were quite robust as the vast majority of them corresponded to a non-parametric *p* value of 0 (i.e., not a single randomly shuffled data set in the permutation process out of 5000 yielded a cluster larger than that which was observed in the actual data). V1, FFA, and pSTS bilaterally all had significantly greater phase locking to both brief and longer exposures within the trial in both α and β frequency bands (Figure 4). The results mirrored those in the time course analyses, in that the phase locking in response to the brief-exposure condition was greater when the stimulus was already off the screen, compared to the longer-exposure condition. The early visual and face-processing regions V1 and FFA were the only regions to show significantly greater phase locking to the longer exposure before the longer-exposure stimulus was removed from the screen (Figure 4). For PAC, we observed significantly greater phase locking to both stimulus exposures for both threat types in the β band. However, activity in the α band was sensitive primarily to longer exposures, with the exception of direct gaze in the left hemisphere, which evoked responses to both the brief and longer-exposure conditions. OFC was engaged primarily by the longer-exposures trials in both bands. The only exception was in left OFC, which displayed stronger β phase locking to the brief exposure during direct-gaze threat cue viewing (*p*=0.05).

**Figure 4.**
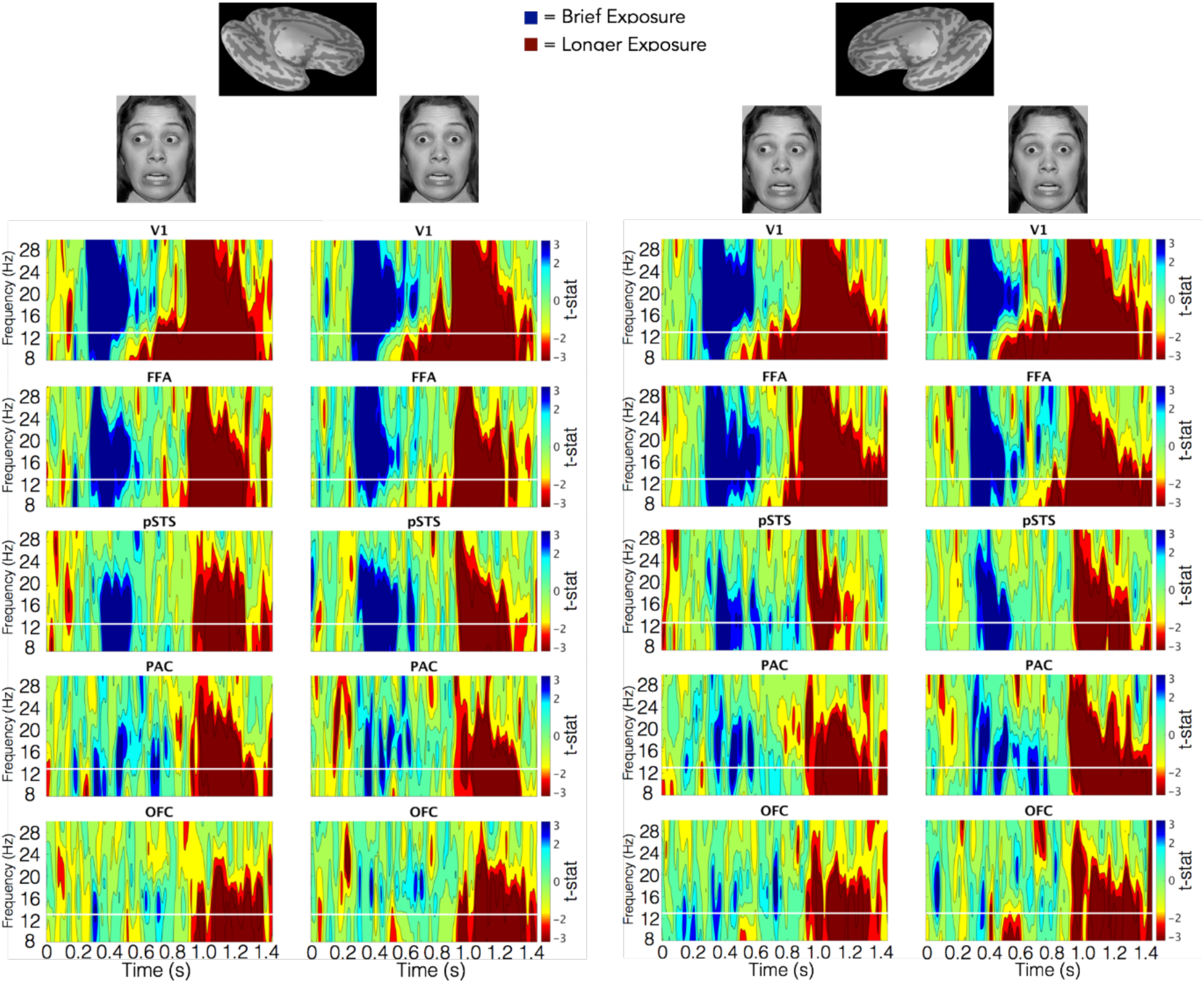
Unthresholded statistical maps of brief-versus longer-exposure phase-locking across trials by threat cue type. The left side shows maps for each ROI from the left hemisphere according to the threat type conveyed by the face at the top of the column (averted-gaze=clear threat, direct-gaze=ambiguous threat). The right side shows the same for the right hemisphere. For each map, the y-axis represents frequency (in Hz) while the x-axis represents time (in s) while the pixel value is the t-statistic representing each participant’s PLF (Phase-locking Factor) across trials for brief exposures (represented in blue) compared to long exposures (represented in red). Contour levels map to significance based on a two-tailed distribution with 59 degrees of freedom. Green represents no significance (i.e. *p-*values above 0.05). The three blue shades (cyan, blue, dark/navy blue) represent p-value ranges between 0.05-0.01,0.01-0.001, and 0.001 and below for brief exposures. The three red shades (yellow, red, dark red) represent the same p-values for long exposures. All *p*-values parametric and uncorrected.

**Table 4.**
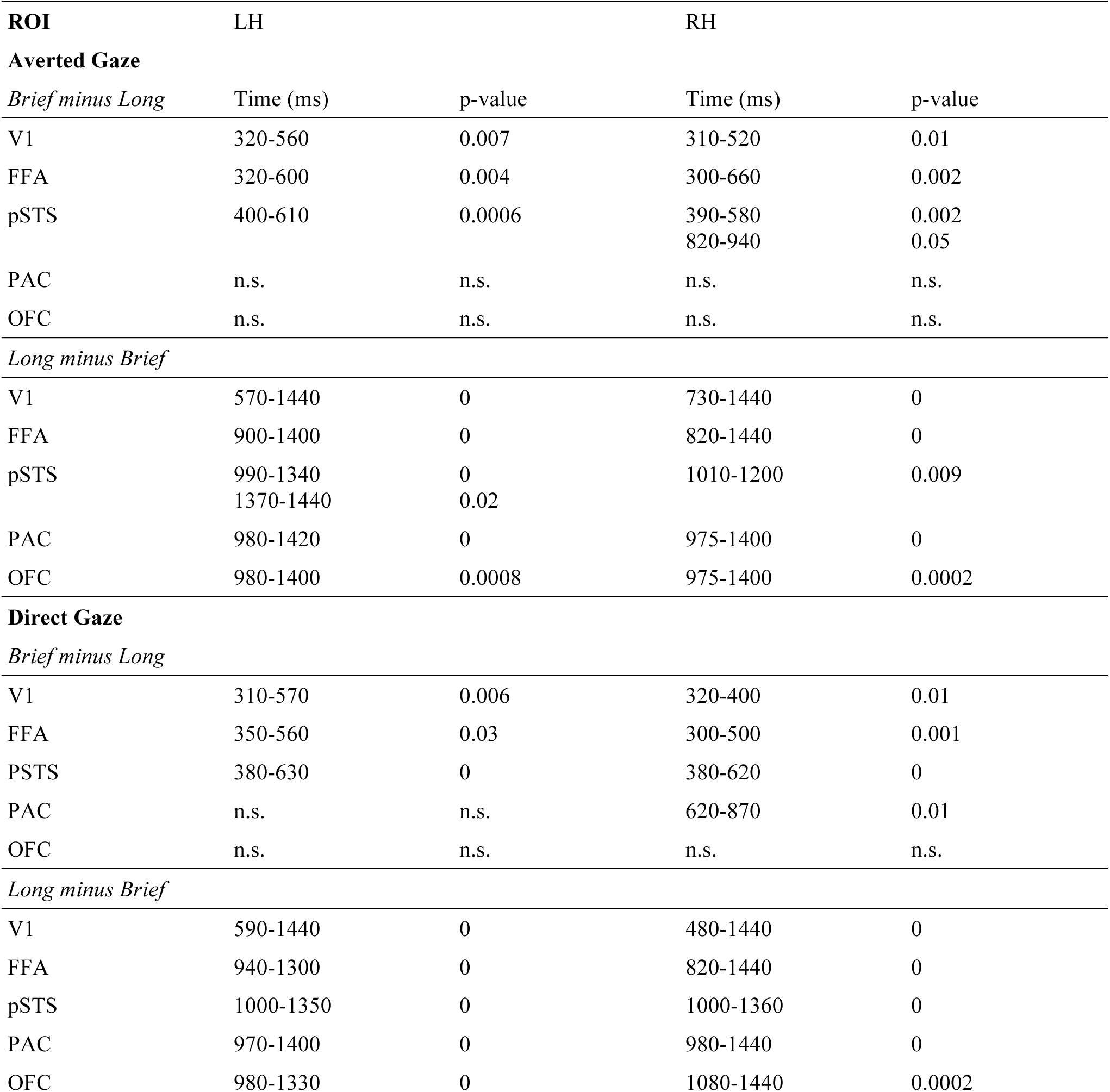
Timing of significant differences in α phase-locking across trials within ROIs between brief (250 ms) and long (883 ms) exposures (reported *p*-values are non-parametric and corrected based on cluster permutations thresholded at *p*<0.05 uncorrected).

| ROI | LH |  | RH |  |
| --- | --- | --- | --- | --- |
| Averted Gaze |  |  |  |  |
| Brief minus Long | Time (ms) | p-value | Time (ms) | p-value |
| V1 | 320-560 | 0.007 | 310-520 | 0.01 |
| FFA | 320-600 | 0.004 | 300-660 | 0.002 |
| pSTS | 400-610 | 0.0006 | 390-580<br>820-940 | 0.002<br>0.05 |
| PAC | n.s. | n.s. | n.s. | n.s. |
| OFC | n.s. | n.s. | n.s. | n.s. |
| Long minus Brief |  |  |  |  |
| V1 | 570-1440 | 0 | 730-1440 | 0 |
| FFA | 900-1400 | 0 | 820-1440 | 0 |
| pSTS | 990-1340<br>1370-1440 | 0<br>0.02 | 1010-1200 | 0.009 |
| PAC | 980-1420 | 0 | 975-1400 | 0 |
| OFC | 980-1400 | 0.0008 | 975-1400 | 0.0002 |
| Direct Gaze |  |  |  |  |
| Brief minus Long |  |  |  |  |
| V1 | 310-570 | 0.006 | 320-400 | 0.01 |
| FFA | 350-560 | 0.03 | 300-500 | 0.001 |
| PSTS | 380-630 | 0 | 380-620 | 0 |
| PAC | n.s. | n.s. | 620-870 | 0.01 |
| OFC | n.s. | n.s. | n.s. | n.s. |
| Long minus Brief |  |  |  |  |
| V1 | 590-1440 | 0 | 480-1440 | 0 |
| FFA | 940-1300 | 0 | 820-1440 | 0 |
| pSTS | 1000-1350 | 0 | 1000-1360 | 0 |
| PAC | 970-1400 | 0 | 980-1440 | 0 |
| OFC | 980-1330 | 0 | 1080-1440 | 0.0002 |

**Table 5.** Timing of significant differences in β phase-locking across trials within ROIs between brief (250 ms) and long (883 ms) exposures (reported *p*-values are non-parametric and corrected based on cluster permutations thresholded at *p*<0.05 uncorrected).

| ROI | LH |  | RH |  |
| --- | --- | --- | --- | --- |
| Averted Gaze |  |  |  |  |
| Brief minus Long | Time (ms) | p-value | Time (ms) | p-value |
| V1 | 300-690 | 0 | 290-650 | 0 |
| FFA | 320-680 | 0 | 320-690 | 0 |
| pSTS | 380-630 | 0 | 390-580 | 0.0002 |
| PAC | 490-700 | 0.005 | 350-570 | 0.002 |
| OFC | n.s. | n.s. | n.s. | n.s. |
| Long minus Brief |  |  |  |  |
| V1 | 750-1400 | 0 | 830-1440 | 0 |
| FFA | 950-1350 | 0 | 920-1440 | 0 |
| pSTS | 980-1330 | 0 | 980-1200 | 0 |
| PAC | 970-1390 | 0 | 980-1200 | 0 |
| OFC | 1100-1420 | 0 | 980-1090<br>1100-1400 | 0.007<br>0 |
| Direct Gaze |  |  |  |  |
| Brief minus Long |  |  |  |  |
| V1 | 310-760 | 0 | 320-660 | 0 |
| FFA | 320-640 | 0 | 330-680<br>760-900 | 0<br>0.04 |
| PSTS | 640-640 | 0 | 360-620 | 0 |
| PAC | 340-460<br>480-710 | 0.01<br>0.0002 | 340-480<br>480-790 | 0.001<br>0.001 |
| OFC | 630-790 | 0.05 | n.s | n.s. |
| Long minus Brief |  |  |  |  |
| V1 | 820-1370 | 0 | 830-1420 | 0 |
| FFA | 970-1300 | 0 | 950-1420 | 0 |
| pSTS | 980-1330 | 0 | 990-1360 | 0 |
| PAC | 970-1390 | 0 | 960-1440 | 0 |
| OFC | 1080-1400 | 0 | 980-1090<br>1090-1440 | 0.003<br>0 |

#### Effects of Gaze: Time Courses

Permutation cluster tests of the time courses within our ROIs revealed only one effect of gaze. We observed significantly increased activation from direct-gaze threat cues during longer exposures in V1 of the right hemisphere late in the trial around 500-650 ms (*p*=0.02). Given its timing, this effect may be the result of feedback from higher regions.

### Effects of Gaze: Phase Locking

#### Brief Stimulus Exposures (250 ms)

##### Significant phase-locking differences for averted-gaze fear faces

The first effect we observed was an increased bilateral response to averted gaze over direct-gaze fear faces. Averted-gaze fear evoked stronger phase locking in the α-band early between left PAC and OFC (*p*=0.013) around 80-160 ms and in right V1 (*p*=0.041) around 140-220 ms (Figure 5A). We also found significantly stronger phase locking for averted-gaze compared to direct-gaze fear in the β-band of right PAC at around 120 ms (*p*=0.023). These findings support our hypothesis of initial reflexive processing being more sensitive to averted-gaze fear, since these congruent cues are thought to be processed more efficiently [15]. In addition, we found stronger phase locking for averted-gaze fear compared to direct-gaze fear very late at the end of the brief-exposure trial. This occurred in the β-band between right FFA and PAC around 1200-1300 ms (*p*=0.012) as well as in the α-band of right PAC from approximately 1320-1380 ms (*p*=0.035) (Figure 5A). Right PAC’s stronger response to averted gaze during brief exposures is in line with our fMRI findings in the right amygdala, which showed greater activation to averted-gaze fear faces compared to direct-gaze fear faces during brief exposures (Figure 5A).

**Figure 5.**
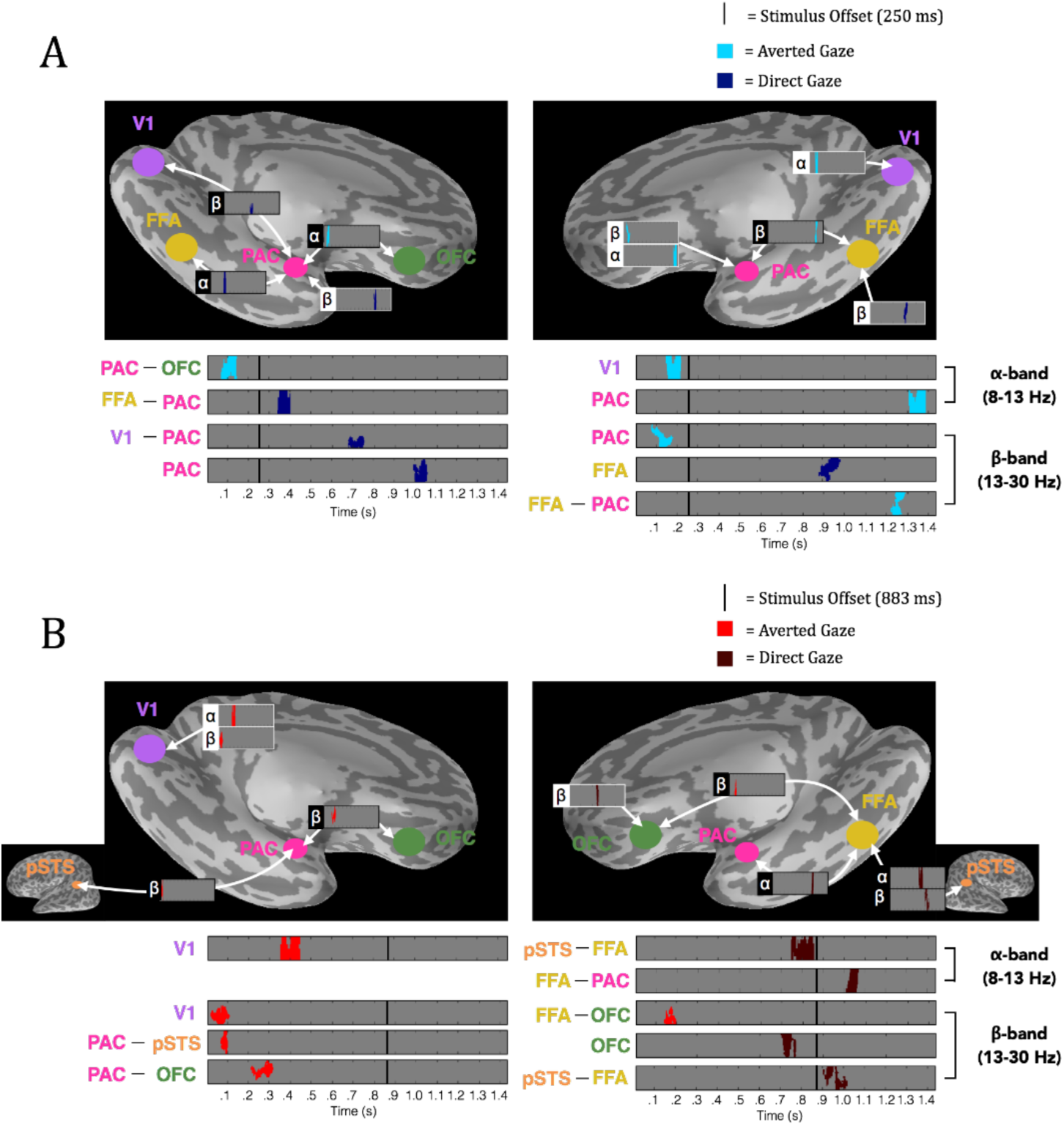
Significant phase-locking for averted vs. direct gaze fearful faces in the extended face-processing network. (***A***) Significant phase-locking for averted-gaze (bright blue) vs. direct-gaze (dark blue) fearful faces during brief exposures (250 ms). **(*B*)** Signifcant phase-locking for averted gaze (bright red) vs. direct gaze (dark red) fearful faces during long exposures (883 ms). *Left* is the left hemisphere phase-locking network (denoted by the image of the left hemisphere). *Right* is the right hemisphere phase-locking network. *Above* is the significant phase-locking overlaid on an inflated *Freesurfer* brain to show anatomical locations of significant phase-locking. White outlines depict phase-locking (PLF) within a region (i.e. phase-locking across trials) while black outlines depict phase-locking values (PLV) between regions. The phase-locking’s frequency band is marked by the appropriate Greek letter to the left (α=8-13Hz, β=13-30 Hz). *Below* is the significant phase-locking magnified and arranged by time of effects as well as frequency band of effects (α on top, β on bottom, as noted on the far right) to display the temporal progression of significant phase-locking through the trial. The anatomical location/connection of the phase-locking is noted to the left. Stimulus offset is marked by the solid black line. For all plots, grey depicts periods of non-significant phase-locking while colored (see legend/A,B descriptions for color to trial type correspondence) blobs depict significant phase-locking determined by non-parametric cluster permutation tests as outlined in *Methods*.

Overall, we found two distinct time windows during the brief-exposure trials in which phase locking, both across trials within a region and in the connectivity between regions, was significantly stronger when exposed to averted-gaze fear faces compared to direct-gaze fear faces: A bilateral response early in the trial around 80-220 ms and then a late response concentrated in the right hemisphere around PAC past 1200 ms (Figure 5A).

##### Significant phase-locking differences for direct-gaze fear faces

For direct-gaze fear faces, there was significantly increased phase locking compared to averted-gaze fear faces during the mid-trial period. We found significantly increased phase locking compared to averted-gaze fear faces in the α-band between left FFA and PAC (*p*=0.04), peaking at around 350 ms. This occurred following the stimulus offset at 250 ms, possibly suggesting that removal of the direct-gaze fear face (incongruent threat cue) elicits sensitivity to this type of threat cue (Figure 5A). Phase locking continued to be stronger to direct-gaze fear in the β-band between left V1 and PAC around 700 ms (*p*=0.001), in the right FFA around 900 ms (*p*=0.05), and in left PAC around 1000 ms (*p*=0.03). Thus, with brief exposures, there is a relative increase of phase locking to direct-gaze fear compared to averted-gaze fear faces beginning immediately after removal of the stimulus up until around 1050 ms.

In summary, during brief exposures we found stronger early *and* late phase locking to averted-gaze fear faces, and stronger phase locking to direct-gaze fear faces mid-trial around the 300-1050 ms range, with right PAC responding more to averted-gaze fear and left PAC responding more to direct-gaze fear. These MEG findings are broadly consistent with the fMRI findings in the amygdalae using this paradigm ([9, 15, 52], Experiment 1 in this study).

#### Longer Stimulus Exposures (883 ms)

##### Significant phase-locking differences for averted-gaze fear faces

With exposures of 883 ms, we likewise observed greater β-band phase locking in response to averted-gaze fear faces early in the trial: in left V1 around 60-80ms (*p*=0.017), between left PAC and pSTS around 80-100ms (*p*=0.032), between right FFA and OFC around 140-200ms (*p*=0.033), and between left PAC and OFC (*p*=0.003) at approximately 200-300 ms (Figure 5B). This again demonstrates a reflexive response more tuned to averted-gaze fear immediately following exposure to the threat cue, suggesting it to be the more salient threat cue to the face processing network. Averted-gaze fear faces also evoked stronger α-band phase locking in left V1 (*p*=0.042) at around 400 ms (Figure 5B) compared to direct-gaze fear. In contrast to brief exposures, stronger phase locking for averted-gaze fear face stimuli persisted until 400 ms suggesting that greater averted-gaze fear phase locking early in the trial is not inherently limited to the pre-300 ms early time frame but can be more sustained when the stimulus exposure is longer. That is, the exposure duration modulates the initial response to averted-gaze fear faces, with longer exposures resulting in longer processing times for averted-gaze fear, when compared to the processing of direct-gaze fear faces under the same exposure conditions.

##### Significant phase-locking differences for direct-gaze fear faces

Towards the latter half of the trial, phase locking became stronger to fearful faces with direct gaze relative to averted gaze in the α-band between pSTS and FFA around 700-950 ms (*p*=0.005) and between FFA and PAC between 1000-1100 ms (*p*=0.034), as shown in Figure 5B. During this same time frame, phase locking in the β-band to direct-gaze fear faces significantly increased over averted-gaze fear faces in right OFC around 700-800 ms (*p*=0.034) and between right pSTS and FFA at roughly 900-1000 ms (*p*=0.01). This provides evidence for our hypothesis that longer presentations result in more extensive processing of direct-gaze fear faces, as all of the significant phase-locking activity differences demonstrated higher phase locking for direct-gaze fear faces late in the trial for connections in the right hemisphere between PAC, FFA, OFC and pSTS.

To conclude, during longer exposures we see similar stronger early phase locking for averted-gaze fear faces relative to direct gaze as we observed in in the brief-exposure trials. However, longer stimulus exposures also prolong this initial processing of averted-gaze fear faces, and delay and prolong processing of direct-gaze fear faces. Returning to greater phase locking for averted-gaze fear faces very late in the trial occurred only during the brief-exposure trials.

## Discussion

The primary aim of the present study was to characterize the neurodynamics mediating facial threat cue perception with MEG, building on previous fMRI findings that employed the same stimulus set of congruent (fearful faces with averted eye gaze) or incongruent (fear with direct gaze) cues [15]. We first validated our experimental design as a direct comparison to previous efforts by replicating the findings of block-design experiments [9, 14, 15] using an event-related paradigm in fMRI. In Experiment 1 (fMRI), we replicated the findings of Adams et al. [15], finding that amygdala responses vary as a function of stimulus exposure duration in response to averted-gaze and direct-gaze fear faces. BOLD activity in the right amygdala was enhanced to averted-gaze fear during brief exposures and direct-gaze fear during longer exposures, whereas the left amygdala activity was greater for longer-exposure durations, particularly for direct-gaze fear. These fMRI findings suggest both that brief stimulus exposures elicit a stronger reflexive response to averted-gaze fear, and that longer exposures evoke more reflective processing of direct-gaze fear, possibly in order to resolve the incongruity in the latter cue combination. These findings align with previous demonstrations of the amygdala being sensitive to brief exposures of averted-gaze fear faces [9, 78] as well as longer exposures of the incongruent threat cues conveyed by direct-gaze fear [14, 79]. However, because of the temporal limitations of fMRI and the BOLD signal, they offer only a partial description of how facial fear cues are processed in the brain, as we detail below.

In Experiment 2 (MEG), we aimed to characterize the effects of exposure duration and compound facial threat cues on the neurodynamics of threat perception on a finer temporal scale by examining the time courses and phase locking of activity in the extended face network, including FFA, PAC (the periamygdaloid cortex), pSTS, and OFC, as well as the primary visual cortex. We observed robust effects of presentation duration in all the ROIs we examined. Brief stimulus exposures for both direct and averted-gaze fear faces resulted in stronger activity immediately following stimulus offset, compared with activity when the stimulus was still present during the longer-exposure trials. In PAC and OFC, this activity persisted for hundreds of milliseconds after the stimulus had been removed in a slow second cycle of processing. Conversely, the longer stimulus exposures elicited significantly greater activation from all our ROIs during the mid-to-late trial period. This would suggest that there is indeed an inherent difference in how these areas respond to facial threat cues, driven by the exposure duration.

We expected brief stimulus presentations to elicit more reflexive threat processing (see also [16–18]), and thus a greater response to averted-gaze fear, which previously had been found to be processed more quickly and efficiently than direct-gaze fear (e.g., [8, 15]). Examination of the phase locking in and between our ROIs indeed revealed a greater initial response to averted-gaze fear in both the α and β frequency ranges. Because of the primary focus on the response in the amygdala complex in previous fMRI studies [9, 14, 15, 52], it should be emphasized that we observed stronger phase locking to averted-gaze fear in the initial response of the right PAC, as well as in the initial connectivity between left PAC and OFC, suggesting more preconscious attention to congruent threat cues in these regions. This finding is consistent with the behavioral findings of Milders et al. [8] and Adams’ shared signal hypothesis [4, 14, 15], suggesting averted-gaze fear is the more salient threat cue. Immediate sensitivity to the more salient threat cue in right PAC is consistent with the right amygdala’s putative function in rapid detection of emotionally salient stimuli [80].

Once the stimulus was removed at 250 ms in the brief presentation condition, we found that phase locking then increased in response to direct-gaze fear faces from about 300-1000 ms, and then was superseded by averted-gaze fear-related activity very late in the trial. Generally, left PAC (along with the left hemisphere as a whole) showed more sensitivity to direct-gaze fear faces during the brief exposures. This is also congruent with the left amygdala’s putative involvement in reflective assessment of stimuli following an initial, reflexive emotional response from the right side [80]. The stronger phase locking to direct-gaze fear during this period reveals a more nuanced picture of PAC activity than that suggested by fMRI results in the amygdala: namely, that the amygdaloid complex appears to be sensitive to both threat cues portrayed by averted and direct-gaze fear at different times. Therefore, the greater BOLD signal evoked by averted-gaze fear during brief exposures in fMRI, as in Experiment 1 and in previous fMRI studies [9, 15] is likely due to the nature of fMRI; that is, summation of (delayed and temporally dilated) BOLD signal, which is unable to resolve fine-grained sensitivity of the amygdala complex to different threat cues at different stages of processing. These results might be partially explained by our finding in MEG that, at the end of the trial (1100-1400 ms), phase locking to congruent threat cues again became stronger than for incongruent threat cues. Significant differences this late in the trial are far too late to be considered “reflexive”, indicating that there is more at play in the neurodynamics mediating threat processing than just an automatic, reflexive response to the brief exposure, followed by a reflective response if stimulus presentation is maintained.

Based on the fMRI findings using this paradigm [9, 14, 15, 52], we had predicted that longer stimulus exposures would evoke a stronger late response to direct-gaze fear, possibly indicative of reflective analysis to resolve ambiguity in these incongruent threat cues. However, because the brief and longer presentation conditions are identical for the first 250 ms, phase locking in and between our ROIs again showed a stronger initial response to averted-gaze fear in both the α and β frequency bands, which resembled the response during the brief stimulus exposures. However, this stronger phase locking to averted gaze was sustained longer, compared with the brief-exposure condition, and extended past the point at which direct-gaze fear-related processing had superseded averted-gaze threat-related processing in the brief duration trials. Similar to brief exposures, mid-trial phase locking during the longer-exposure trials was stronger to direct-gaze fear faces compared to averted-gaze fear faces, but this sensitivity to threat signal incongruity began later and was not superseded by increased phase locking to congruent threat cues later in the trial, as in the brief-exposure trials.

In addition to the temporal differences, we found differences in the spatial pattern of significant phase locking. Compared to the phase locking during brief-exposure trials, which was concentrated around left PAC, phase locking during longer exposures was centered mostly around right FFA. Reflective processing is associated more with the ventral stream [16], of which the FFA is a key module. Thus, greater involvement of FFA during longer-exposure trials may provide some support to our hypothesis of reflective processing being biased toward ventral stream processing. A notable connection for right FFA during longer exposures was to right pSTS, which displayed two temporally and spectrally distinct periods of incongruent threat-cue sensitivity immediately preceding and following stimulus offset. The late involvement of pSTS during longer exposures could potentially be representative of its function as part of the “social brain” [81], as the right pSTS in particular has been linked to internalizing another’s perspective [82]. It is thought to be a key region for visual integration of social cues, especially when inferring social meaning from combinations of facial expression and eye gaze [83], making it a sensible candidate to be recruited in resolving possible ambiguity of the incongruent threat cue during longer exposures. Collectively, the phase locking seen in the right hemisphere supports the idea of slower reflective analysis being mediated by the ‘high road’ [78] during longer exposures to resolve the meaning of the incongruent threat cue. This appears at odds with the left hemisphere’s role as being primarily responsible for reflective analysis in previous findings [84]. One speculative resolution of this incongruity could lie in the general right hemisphere specialization for face processing [85, 86] as well as in the right STS’s specialization in gaze perception [87]. The nature of such late activity is difficult to interpret, however, particularly since most studies on the topic have been done with fMRI and not explicitly designed to test the lateralization of the amygdala responses.

In our phase-locking results, we did not see much evidence for functional separation between the α and β bands. However, compared to other frequency bands, α-band activity does not fit into a general hypothesis of function [88, 89]. In the specific context of α activity evoked by viewing emotional faces, α responses have been shown to be higher in response to negative (angry) compared to positive (happy) emotional faces over temporal, parietal, and occipital recording sites [90], but weaker in response to valenced faces (both positive and negative) compared to neutral faces [91, 92]. The evidence for any specific α-band function in emotional perception is sparse and inconsistent, but none of it suggests the role of inhibition [89]. We found α-band activity to be sensitive to negative valence, as α phase locking was stronger both in initial reflexive processing of averted-gaze fear, suggesting a role in increasing shared-signal processing efficiency, and in late reflective processing of the direct-gaze fear (incongruent signal). In addition, α-phase locking was mostly congruent with β activity both in effects of stimulus duration and effects of eye-gaze, supporting an active non-inhibitory role for α-band oscillations during facial threat cue perception, at least in terms of its phase locking (compared to power or other magnitude-dependent measures).

In summary, with fMRI, we replicated previous stimulus exposure duration effects for fear expression processing as a function of gaze perception using a broad stimulus set and employing an event-related design, and showing that previous findings using this paradigm [9, 14, 15, 52] are not dependent upon states induced by a block design. This set the stage for using this identical paradigm in MEG to examine the temporal dynamics of the effect. We found that the stimulus exposure duration strongly modulated the fine-grained neurodynamics in the extended face perception network, resulting in a slower processing sequence for the longer-exposure stimuli. However, the effects of gaze, while modulated by the exposure duration, were nonetheless evident in greater early phase locking to averted gaze, providing neural evidence at a high temporal resolution that these congruent compound cues are indeed allocated more preconscious attention, and in increased later phase locking to direct gaze, again supporting the idea that slow, reflective processing is biased towards the incongruent threat cue. However, the fast neurodynamics in the extended face-processing network in response to these brief and longer exposures observed with MEG show that its response is far more complicated than a simple dichotomy of a reflexive response induced by a brief stimulus exposure and a reflective response induced by a longer stimulus presentation. Here we have shown that the observed greater BOLD response to averted-gaze fear in the amygdala complex during brief exposures is not the result of the processing being curtailed by the brief presentation, and thus cutting off reflective processing. Rather, it likely stems from the summation of the hemodynamic signal over the entire trial in fMRI, as we observed stronger mid-trial phase locking to direct-gaze fear during both brief and longer exposures, and the initial reflexive response to averted-gaze fear, again regardless of exposure duration. Previous attempts to rationalize the opposing findings in fMRI suggested that congruent cues such as averted-gaze fear not only capture attention more readily, but due to their aversive nature, may result in more rapid disengagement, explaining the comparative dwelling of the neural response on direct-gaze fear faces during longer exposures [93]. The ability to empirically address this speculation with the high temporal resolution of MEG revealed that this does not in fact seem to be the case, as we observed prolonged cognitive engagement with averted-gaze fear relative to direct-gaze fear under longer exposures. Our findings thus underscore the importance of investigating brain activity using techniques with high temporal resolution to gain a more complete picture of the neurodynamics underlying perception of compound threat cues from the face.

## Acknowledgements

This work was supported with the following funds:

NIMH R01MH101194 awarded to KK and RBA, Jr.

NIMH K01MH084011 awarded to KK

NCRR P41RR14075 to A.A. Martinos Center

Instrumentation Grants 1S10RR023401, 1S10RR019307, and 1S10RR023043.

## Author Contributions

CC contributed data collection, analysis, and writing the paper.

HY contributed data collection, analysis, and writing the paper.

RBA, Jr. contributed funding, study design/conception, and writing the paper.

NW, DNA, and TGS contributed data collection and manuscript editing.

KK contributed funding, study design/conception, data collection, analysis, and writing the paper.

## References

1. Adams, R. B., Ambady, N., Nakayama, K., & Shimojo, S. (2011). The Science of Social Vision. The Science of Social Vision. doi:10.1093/acprof:oso/9780195333176.001.0001

2. Adams, R. B., Ambady, N., Macrae, C. N., & Kleck, R. E. (2006). Emotional expressions forecast approach-avoidance behavior. Motivation and Emotion, 30 (2), 179–188. doi:10.1007/s 11031-006-9020-2

3. Adams, R. B., & Kleck, R. E. (2003). Perceived Gaze direction and the processing of facial displays of emotion. Psychological Science, 14(6), 644–647. doi:10.1046/j.0956-7976.2003.psci_1479.x

4. Adams, R. B. J., & Kleck, R. E. (2005). Effects of Direct and Averted Gaze on the Perception of Facially Communicated Emotion. Emotion, 5(1), 3–11. doi:10.1037/1528-3542.5.1.3

5. Sander, D., Grandjean, D., Kaiser, S., Wehrle, T., & Scherer, K. R. (2007). Interaction effects of perceived gaze direction and dynamic facial expression: Evidence for appraisal theories of emotion. European Journal of Cognitive Psychology, 19(3), 470–480. doi:10.1080/09541440600757426

6. Benton, C. P. (2010). Rapid reactions to direct and averted facial expressions of fear and anger. Visual Cognition, 18(9), 1298–1319. doi:10.1080/13506285.2010.481874

7. Fox, E., Mathews, A., Calder, A. J., & Yiend, J. (2007). Anxiety and sensitivity to gaze direction in emotionally expressive faces. Emotion (Washington, D.C.), 7(3), 478–486. doi:10.1037/1528- 3542.7.3.478

8. Milders, M., Hietanen, J. K., Leppänen, J. M., & Braun, M. (2011). Detection of emotional faces is modulated by the direction of eye gaze. Emotion, 11(6), 1456–1461. doi:10.1037/a0022901

9. Hadjikhani, N., Hoge, R., Snyder, J., & de Gelder, B. (2008). Pointing with the eyes: The role of gaze in communicating danger. Brain and Cognition, 68(1), 1–8. doi:10.1016/j.bandc.2008.01.008

10. N’Diaye, K., Sander, D., & Vuilleumier, P. (2009). Self-relevance processing in the human amygdala: gaze direction, facial expression, and emotion intensity. Emotion, 9(6), 798–806. doi:10.1037/a0017845

11. Sato, W., Yoshikawa, S., Kochiyama, T., & Matsumura, M. (2004). The amygdala processes the emotional significance of facial expressions: An fMRI investigation using the interaction between expression and face direction. NeuroImage, 22(2), 1006–1013. doi:10.101 6/j.neuroimage.2004.02.030

12. Whalen, P. J., Shin, L. M., McInerney, S. C., Fischer, H., Wright, C. I., & Rauch, S. L. (2001). A functional MRI study of human amygdala responses to facial expressions of fear versus anger. Emotion, 1(1), 70–83. doi:10.1037/1528-3542.1.1.70

13. Adams, R. B., & Janata, P. (2002). A comparison of neural circuits underlying auditory and visual object categorization. NeuroImage, 16, 361–377. doi:10.1006/nimg.2002.1088

14. Adams Jr., R. B., Gordon, H. L., Baird, A. A., Ambady, N., & Kleck, R. E. (2003). Effects of gaze on amygdala sensitivity to anger and fear faces. Science, 300(5625), 1536. doi:10.1 126/science. 1082244

15. Adams, R. B., Franklin, R. G., Kveraga, K., Ambady, N., Kleck, R. E., Whalen, P. J., … Nelson, A. J. (2012). Amygdala responses to averted vs direct gaze fear vary as a function of presentation speed. Social Cognitive and Affective Neuroscience, 7(5), 568–577. doi:10.1093/scan/nsr038

16. Lieberman, M. D., Gaunt, R., Gilbert, D. T., & Trope, Y. (2002). Reflexion and reflection: A social cognitive neuroscience approach to attributional inference. Advances in Experimental Social Psychology, 34, 199–249. doi:10.1016/S0065-2601(02)80006-5

17. Lieberman, M. D. (2003). Reflexive and reflective judgment processes: A social cognitive neuroscience approach. Social judgments: Implicit and explicit processes, 44–67. doi:10.1037//0033-2909.126.1.109

18. Cunningham, W. A., & Zelazo, P. D. (2007). Attitudes and evaluations: a social cognitive neuroscience perspective. Trends in Cognitive Sciences, 11(3), 97–104. doi:10.1016/j.tics.2006.12.005

19. Ewbank, M. P., Fox, E., & Calder, A. J. (2010). The Interaction Between Gaze and Facial Expression in the Amygdala and Extended Amygdala is Modulated by Anxiety. Frontiers in human neuroscience, 4, 56. doi:10.3389/fnhum.2010.00056

20. Wicker, B., Michel, F., Henaff, M. a, & Decety, J. (1998). Brain regions involved in the perception of gaze: a PET study. NeuroImage, 8(2), 221–7. doi:10.1006/nimg.1998.0357

21. Hooker, C. I., Paller, K. A., Gitelman, D. R., Parrish, T. B., Mesulam, M. M., & Reber, P. J. (2003). Brain networks for analyzing eye gaze. Cognitive Brain Research, 17(2), 406–418. doi:10.1016/S0926-6410(03)00143-5

22. Haxby, J. V., Hoffman, E. A., & Gobbini, M. I. (2002). Human neural systems for face recognition and social communication. Biological Psychiatry, 51(1), 59–67. doi:10.1016/S0006-3223(01)01330-0

23. Hardee, J. E., Thompson, J. C. J. C., & Puce, A. (2008). The left amygdala knows fear: Laterality in the amygdala response to fearful eyes. Social Cognitive and Affective Neuroscience, 3(1), 47–54. doi:10.1093/scan/nsn001

24. Puce, A., Allison, T., Gore, J. C., & McCarthy, G. (1995). Face-sensitive regions in human extrastriate cortex studied by functional MRI. Journal of Neurophysiology, 74(3), 1192–9. Retrieved from http://www.ncbi.nlm.nih.gov/entrez/query.fcgi?cmd=Retrieve&db=PubMed&dopt=Citation&list_uids=7500143

25. Hoffman, E. a, & Haxby, J. V. (2000). Distinct representations of eye gaze and identity in the distributed human neural system for face perception. Nature neuroscience, 3(1), 80–84. doi:10.1038/71152

26. George, N., Driver, J., & Dolan, R. J. (2001). Seen gaze-direction modulates fusiform activity and its coupling with other brain areas during face processing. NeuroImage, 13(6 Pt 1), 1102–1112. doi:10.1006/nimg.2001.0769

27. Harris, R. J., Young, A. W., & Andrews, T. J. (2012). Morphing between expressions dissociates continuous from categorical representations of facial expression in the human brain. Proceedings of the National Academy of Sciences of the United States of America, 109, 21164–21169. doi:10.1073/pnas.1212207110/-

28. Baron-Cohen, S., Baron-Cohen, S., Ring, H. A., Ring, H. A., Wheelright, S., Wheelright, S., … Williams, S. C. (1999). Social intelligence in the normal and autisitic brain: an fMRI study. European Journal of Neuroscience, 11(6), 1891–1898. doi:10.1046/j.1460-9568.1999.00621.x

29. Nummenmaa, L., & Calder, A. J. (2009). Neural mechanisms of social attention. Trends in Cognitive Sciences. doi:10.1016/j.tics.2008.12.006

30. Rhodes, G., Calder, A., Johnson, M., & Haxby, J. V. (2012). Oxford Handbook of Face Perception. Oxford Handbook of Face Perception. doi:10.1093/oxfordhb/9780199559053.001.0001

31. Attal, Y., Bhattacharjee, M., Yelnik, J., Cottereau, B., Lefèvre, J., Okada, Y., … Baillet, S. (2007). Modeling and detecting deep brain activity with MEG & EEG. In Annual International Conference of the IEEE Engineering in Medicine and Biology - Proceedings (pp. 4937–4940). doi:10.11 09/IEMBS.2007.4353448

32. Dumas, T., Attal, Y., Chupin, M., Jouvent, R., Dubal, S., & George, N. (2010). MEG study of amygdala responses during the perception of emotional faces and gaze. In IFMBE Proceedings (Vol. 28, pp. 330–333). doi:10.1007/978-3-642-12197-5_77

33. Attal, Y., & Schwartz, D. (2013). Assessment of Subcortical Source Localization Using Deep Brain Activity Imaging Model with Minimum Norm Operators: A MEG Study. PLoS ONE, 8(3). doi:10.1371/journal.pone.0059856

34. Dumas, T., Dubal, S., Attal, Y., Chupin, M., Jouvent, R., Morel, S., & George, N. (2013). MEG Evidence for Dynamic Amygdala Modulations by Gaze and Facial Emotions. PLoS ONE, 8(9). doi:10.1371/journal.pone.0074145

35. Dumas, T., Attal, Y., Dubal, S., Jouvent, R., & George, N. (2011). Detection of activity from the amygdala with magnetoencephalography. IRBM, 32(1), 42–47. doi:10.1016/j.irbm.2010.11.001

36. Aggleton, J. P. (2000). The Amygdala: a functional analysis. Oxford University Press, USA. Retrieved from http://orca.cf.ac.uk/34922/

37. Pitkänen, a, Pikkarainen, M., Nurminen, N., & Ylinen, a. (2000). Reciprocal connections between the amygdala and the hippocampal formation, perirhinal cortex, and postrhinal cortex in rat. A review. Annals of the New York Academy of Sciences, 911, 369–391. doi:10.1111/j.1749- 6632.2000.tb06738.x

38. Klimesch, W., Freunberger, R., Sauseng, P., & Gruber, W. (2008). A short review of slow phase synchronization and memory: Evidence for control processes in different memory systems? Brain Research. doi:10.1016/j.brainres.2008.06.049

39. Düzel, E., Neufang, M., & Heinze, H. J. (2005). The oscillatory dynamics of recognition memory and its relationship to event-related responses. Cerebral Cortex, 15(12), 1992–2002. doi:10.1093/cercor/bhi074

40. Banerjee, S., Snyder, A. C., Molholm, S., & Foxe, J. J. (2011). Oscillatory alpha-band mechanisms and the deployment of spatial attention to anticipated auditory and visual target locations: supramodal or sensory-specific control mechanisms? The Journal of neuroscience: the official journal of the Society for Neuroscience, 31(27), 9923–9932. doi:10.1523/JNEUROSCI.4660-10.2011

41. Haegens, S., Nácher, V., Luna, R., Romo, R., & Jensen, O. (2011). α-Oscillations in the monkey sensorimotor network influence discrimination performance by rhythmical inhibition of neuronal spiking. Proceedings of the National Academy of Sciences of the United States of America, 108(48), 19377–82. doi:10.1073/pnas.1117190108

42. Romei, V., Gross, J., & Thut, G. (2010). On the role of prestimulus alpha rhythms over occipitoparietal areas in visual input regulation: correlation or causation? The Journal of neuroscience: the official journal of the Society for Neuroscience, 30(25), 8692–8697. doi:10.1523/JNEUROSCI.0160-10.2010

43. Thut, G. (2006). -Band Electroencephalographic Activity over Occipital Cortex Indexes Visuospatial Attention Bias and Predicts Visual Target Detection. Journal of Neuroscience, 26(37), 9494–9502. doi:10.1523/JNEUROSCI.0875-06.2006

44. Kveraga, K., Ghuman, A. S., Kassam, K. S., Aminoff, E. A., Hämäläinen, M. S., Chaumon, M., & Bar, M. (2011). Early onset of neural synchronization in the contextual associations network. Proceedings of the National Academy of Sciences of the United States of America, 108(8), 3389–3394. doi:10.1073/pnas.1013760108

45. Okazaki, M., Kaneko, Y., Yumoto, M., & Arima, K. (2008). Perceptual change in response to a bistable picture increases neuromagnetic beta-band activities. Neuroscience Research, 61(3), 319–328. doi:10.1016/j.neures.2008.03.010

46. Sehatpour, P., Molholm, S., Schwartz, T. H., Mahoney, J. R., Mehta, A. D., Javitt, D. C., … Foxe, J. J. (2008). A human intracranial study of long-range oscillatory coherence across a frontal-occipital-hippocampal brain network during visual object processing. Proceedings of the National Academy of Sciences of the United States of America, 105(11), 4399–4404. doi:10.1073/pnas.0708418105

47. Senkowski, D., Molholm, S., Gomez-Ramirez, M., & Foxe, J. J. (2006). Oscillatory beta activity predicts response speed during a multisensory audiovisual reaction time task: A high-density electrical mapping study. Cerebral Cortex, 16(11), 1556–1565. doi:10.1093/cercor/bhj091

48. Posner, M. I. (1980). Orienting of attention. Q J Exp Psychol, 32(September), 3–25. doi:10.1080/00335558008248231

49. Driver, J., Davis, G., Ricciardelli, P., Kidd, P., Maxwell, E., & Baron-Cohen, S. (1999). Gaze perception triggers reflexive visuospatial orienting. Visual Cognition, 6(5), 509–540. doi:10.1080/ 135062899394920

50. Cooper, R. M., & Langton, S. R. H. (2006). Attentional bias to angry faces using the dot-probe task? It depends when you look for it. Behaviour Research and Therapy, 44(9), 1321–1329. doi:10.1016/j.brat.2005.10.004

51. Jones, B. C., DeBruine, L. M., Main, J. C., Little, A. C., Welling, L. L. M., Feinberg, D. R., & Tiddeman, B. P. (2010). Facial cues of dominance modulate the short-term gaze-cuing effect in human observers. Proceedings. Biological sciences / The Royal Society, 277(1681), 617–624. doi:10.1098/rspb.2009.1575

52. Van Der Zwaag, W., Da Costa, S. E., Zürcher, N. R., Adams, R. B., & Hadjikhani, N. (2012). A 7 tesla fMRI study of amygdala responses to fearful faces. Brain Topography, 25(2), 125–128. doi:10.1007/s10548-012-0219-0

53. Sharon, D., Hämäläinen, M. S., Tootell, R. B. H., Halgren, E., & Belliveau, J. W. (2007). The advantage of combining MEG and EEG: Comparison to fMRI in focally stimulated visual cortex. NeuroImage, 36(4), 1225–1235. doi:10.1016/j.neuroimage.2007.03.066

54. Ekman, R. (1975). Pictures of Facial Affect. Emotion, 1–8. doi:citeulike-article-id:4270156

55. Tottenham, N., Tanaka, J. W., Leon, A. C., McCarry, T., Nurse, M., Hare, T. a., … Nelson, C. (2009). The NimStim set of facial expressions: Judgements from untrained research participants. Psychiatry Research, 168(3), 242–249. doi:10.1016/j.psychres.2008.05.006. The

56. Brainard, D. H. (1997). The Psychophysics Toolbox. Spatial Vision, 10, 433–436. doi:10.1163/156856897X00357

57. Pelli, D. G. (1997). The VideoToolbox software for visual psychophysics: transforming numbers into movies. Spatial Vision. doi:10.1163/156856897X00366

58. Lieberman, M., Inagaki, T. K., Tabibnia, G., & Crockett, M. J. (2011). Subjective responses to emotional stimuli during labeling, reappraisal, and distraction. Emotoin, 11(3), 468–480. doi:10.1037/a0023503

59. Deichmann, R., Gottfried, J. A., Hutton, C., & Turner, R. (2003). Optimized EPI for fMRI studies of the orbitofrontal cortex. NeuroImage, 19(2), 430–441. doi:10.1016/S1053- 8119(03)00073-9

60. Kveraga, K., Boshyan, J., & Bar, M. (2007). Magnocellular projections as the trigger of top-down facilitation in recognition. The Journal of neuroscience: the official journal of the Society for Neuroscience, 27(48), 13232–40. doi:10.1523/JNEUROSCI.3481-07.2007

61. Kringelbach, M. L., & Rolls, E. T. (2004). The functional neuroanatomy of the human orbitofrontal cortex: Evidence from neuroimaging and neuropsychology. Progress in Neurobiology. doi:10.1016/j.pneurobio.2004.03.006

62. Mazaika, Hoeft, Glovera, Mazaika, P. K., Hoeft, F., Glover, G. H., & Reiss, A. L. (2009). Methods and Software for fMRI Analysis for Clinical Subjects. Human Brain Mapping, 77309. doi:10.1016/S 1053-8119(09)70238-1

63. Gramfort, A., Luessi, M., Larson, E., Engemann, D. A., Strohmeier, D., Brodbeck, C., … Hämäläinen, M. S. (2014). MNE software for processing MEG and EEG data. NeuroImage, 86, 446–460. doi:10.1016/j.neuroimage.2013.10.027

64. Gramfort, A., Luessi, M., Larson, E., Engemann, D. A., Strohmeier, D., Brodbeck, C., … Hämäläinen, M. (2013). MEG and EEG data analysis with MNE-Python. Frontiers in Neuroscience, (7 DEC). doi:10.3389/fnins.2013.00267

65. Tesche, C. D., Uusitalo, M. A., Ilmoniemi, R. J., Huotilainen, M., Kajola, M., & Salonen, O. (1995). Signal-space projections of MEG data characterize both distributed and well-localized neuronal sources. Electroencephalography and Clinical Neurophysiology, 95(3), 189–200. doi:10.1016/0013-4694(95)00064-6

66. Uusitalo, M. a, & Ilmoniemi, R. J. (1997). Signal-space projection method for separating MEG or EEG into components. Medical & biological engineering & computing, 35(2), 135–140. doi:10.1007/BF02534144

67. Fischl, B., Salat, D. H., Van Der Kouwe, A. J. W., Makris, N., Ségonne, F., Quinn, B. T., & Dale, A. M. (2004). Sequence-independent segmentation of magnetic resonance images. In NeuroImage (Vol. 23). doi:10.1016/j.neuroimage.2004.07.016

68. Lin, F. H., Belliveau, J. W., Dale, A. M., & Hämäläinen, M. S. (2006). Distributed current estimates using cortical orientation constraints. Human Brain Mapping, 27(1), 1–13. doi:10.1002/hbm.20155

69. Dale, A. M., Liu, A. K., Fischl, B. R., Buckner, R. L., Belliveau, J. W., Lewine, J. D., & Halgren, E. (2000). Dynamic Statistical Parametric Mapping: Combining fMRI and MEG for High-Resolution Imaging of Cortical Activity. Neuron, 26(1), 55–67. doi:10.1016/S0896- 6273(00)81138-1

70. Engemann, D. A., & Gramfort, A. (2015). Automated model selection in covariance estimation and spatial whitening of MEG and EEG signals. NeuroImage, 108, 328–342. doi:10.1016/j.neuroimage.2014.12.040

71. Lachaux, J. P., Rodriguez, E., Martinerie, J., & Varela, F. J. (1999). Measuring phase synchrony in brain signals. Human Brain Mapping, 8(4), 194–208. doi:10.1002/(SICI)1097-0193(1999)8:4<194::AID-HBM4>3.0.CO;2-C

72. Maris, E., & Oostenveld, R. (2007). Nonparametric statistical testing of EEG- and MEG-data. Journal of Neuroscience Methods, 164(1), 177–190. doi:10.1016/j.jneumeth.2007.03.024

73. Adams, R. B., Franklin, R. G., Nelson, A. J., Gordon, H. L., Kleck, R. E., Whalen, P. J., & Ambady, N. (2011). Differentially tuned responses to restricted versus prolonged awareness of threat: A preliminary fMRI investigation. Brain and Cognition, 77(1), 113–119. doi:10.1016/j.bandc.2011.05.001

74. Dien, J. (2009). A tale of two recognition systems: Implications of the fusiform face area and the visual word form area for lateralized object recognition models. Neuropsychologia, 47(1), 1–16. doi:10.1016/j.neuropsychologia.2008.08.024

75. Hamilton, C. R., & Vermeire, B. a. (1988). Complementary hemispheric specialization in monkeys. Science (New York, N.Y.), 242(4886), 1691–4. doi:10.1111/j.1749- 6632.1977.tb41909.x

76. Kanwisher, N., McDermott, J., & Chun, M. M. (1997). The fusiform face area: a module in human extrastriate cortex specialized for face perception. The Journal of neuroscience: the official journal of the Society for Neuroscience, 17(11), 4302–11. doi:10.1098/Rstb.2006.1934

77. Yovel, G., Tambini, A., & Brandman, T. (2008). The asymmetry of the fusiform face area is a stable individual characteristic that underlies the left-visual-field superiority for faces. Neuropsychologia, 46(13), 3061–3068. doi:10.1016/j.neuropsychologia.2008.06.017

78. LeDoux, J. E. (1998). The emotional brain: The mysterious underpinnings of emotional life. The emotional brain: The mysterious underpinnings of emotional life. doi:10.2307/3953278

79. Hsu, M., Bhatt, M., Adolphs, R., Tranel, D., & Camerer, C. F. (2005). Neural systems responding to degrees of uncertainty in human decision-making. Science (New York, N.Y.), 310(5754), 1680–3. doi:10.1126/science.1115327

80. Rogers, L. J., Vallortigara, G., & Andrew, R. J. (2013). Divided Brains. Divided Brains: The Biology and Behaviour of Brain Asymmetries. doi:10.1017/CBO9780511793899

81. Lahnakoski, J. M., Glerean, E., Salmi, J., Jääskeläinen, I. P., Sams, M., Hari, R., & Nummenmaa, L. (2012). Naturalistic FMRI mapping reveals superior temporal sulcus as the hub for the distributed brain network for social perception. Frontiers in human neuroscience, 6(August), 233. doi:10.3389/fnhum.2012.00233

82. David, N., Aumann, C., Santos, N. S., Bewernick, B. H., Eickhoff, S. B., Newen, A., … Vogeley, K. (2008). Differential involvement of the posterior temporal cortex in mentalizing but not perspective taking. Social Cognitive and Affective Neuroscience, 3(3), 279–289. doi:10.1093/scan/nsn023

83. Adams, R. B., Rule, N. O., Franklin, R. G., Wang, E., Stevenson, M. T., Yoshikawa, S., … Ambady, N. (2010). Cross-cultural reading the mind in the eyes: an fMRI investigation. Journal of Cognitive Neuroscience, 22(1), 97–108. doi:10.1162/jocn.2009.21187

84. Morris, J. S., Ohman, A., & Dolan, R. J. (1998). Conscious and unconscious emotional learning in the human amygdala. Nature, 393(6684), 467–470. doi:10.1038/30976

85. Zangenehpour, S., & Chaudhuri, A. (2005). Patchy organization and asymmetric distribution of the neural correlates of face processing in monkey inferotemporal cortex. Current Biology, 15(11), 993–1005. doi:10.1016/j.cub.2005.04.031

86. Tsao, D. Y., & Livingstone, M. S. (2008). Mechanisms of face perception. Annual review of neuroscience, 31, 411–437. doi:10.1146/annurev.neuro.30.051606.094238.Mechanisms

87. Calder, A. J., Beaver, J. D., Winston, J. S., Dolan, R. J., Jenkins, R., Eger, E., & Henson, R. N. A. (2007). Separate Coding of Different Gaze Directions in the Superior Temporal Sulcus and Inferior Parietal Lobule. Current Biology, 17(1), 20–25. doi:10.1016/j.cub.2006.10.052

88. Bazar, E., & Güntekin, B. (2012). A short review of alpha activity in cognitive processes and in cognitive impairment. International Journal of Psychophysiology. doi:10.1016/j.ijpsycho.2012.07.001

89. Güntekin, B., & Başar, E. (2014). A review of brain oscillations in perception of faces and emotional pictures. Neuropsychologia. doi:10.1016/j.neuropsychologia.2014.03.014

90. Güntekin, B., & Basar, E. (2007). Emotional face expressions are differentiated with brain oscillations. International Journal of Psychophysiology, 64(1), 91–100.doi:10.1016/j.ijpsycho.2006.07.003

91. Balconi, M., & Mazza, G. (2009). Brain oscillations and BIS/BAS (behavioral inhibition/activation system) effects on processing masked emotional cues. ERS/ERD and coherence measures of alpha band. International Journal of Psychophysiology, 74(2), 158–165. doi:10.1016/j.ijpsycho.2009.08.006

92. Balconi, M., Brambilla, E., & Falbo, L. (2009). BIS/BAS, cortical oscillations and coherence in response to emotional cues. Brain Research Bulletin, 80(3), 151–157. doi:10.1016/j.brainresbull.2009.07.001

93. Mogg, K., Garner, M., & Bradley, B. P. (2007). Anxiety and orienting of gaze to angry and fearful faces. Biological Psychology, 76(3), 163–169. doi:10.1016/j.biopsycho.2007.07.005

